# Seasonal bee communities vary in their responses to local and landscape scales: implication for land managers

**DOI:** 10.1101/2022.07.13.499741

**Authors:** Melanie Kammerer, Aaron L. Iverson, Kevin Li, John F. Tooker, Christina M. Grozinger

## Abstract

**Context:** There is great interest in land management practices for pollinators; however, a quantitative comparison of landscape and local effects on bee communities is necessary to determine if adding small habitat patches can increase bee abundance or species richness. The value of increasing floral abundance at a site is undoubtedly influenced by the phenology and magnitude of floral resources in the landscape, but due to the complexity of measuring landscape-scale resources, these factors have been understudied.

**Objectives:** To address this knowledge gap, we quantified the relative importance of local versus landscape scale resources for bee communities, identified the most important metrics of local and landscape quality, and evaluated how these relationships vary with season.

**Methods:** We studied season-specific relationships between local and landscape quality and wild-bee communities at 33 sites in the Finger Lakes region of New York, USA. We paired site surveys of wild bees, plants, and soil characteristics with a multi-dimensional assessment of landscape composition, configuration, insecticide toxic load, and a spatio-temporal evaluation of floral resources at local and landscape scales.

**Results:** We found that the most relevant spatial scale varied by season. Spring bees depended on landscape resources, but summer bees responded more to local quality, implying that site-level management is most likely to be successful in supporting summer bees. Semi-natural habitats, including forests, wetlands, and other aquatic habitats, were particularly important for spring bees.

**Conclusions:** By considering spatial and temporal variation in resources, we developed season-specific recommendations to improve habitat quality for wild bees and offset manifold stressors threatening these essential pollinators.

## 1. Introduction

There is great interest in developing approaches for translational ecology (Schlesinger 2010; Enquist et al. 2017), where research is designed to provide stakeholders with information they can use to address challenges. Because of the importance of pollinators for agricultural production and resilience of ecological communities, there has been substantial investment in land management practices to increase abundance and diversity of pollinator populations. Indeed, from 2006-2016, the United States and Europe spent more than 35 billion USD on agri-environment schemes (Ansell et al. 2016), including practices targeting pollinators. While most conservation practices tend to be implemented at local scales, many recent studies of pollinator ecology focus on landscape-level phenomena, creating a disconnect with the scale at which conservation is implemented. In addition, despite documented seasonal shifts in pollinator communities and the floral resources on which they depend for nutrition, temporally resolved information on pollinator and floral communities is typically not considered. To best inform agri-environment schemes, it is necessary to understand how local and landscape quality, across seasons, co-influence pollinator communities.

Conservation practices typically ameliorate very small proportions of landscapes that pollinators experience. The spatial extent of a landscape should be ‘organism-defined’ (Wiens 1976), which for wild bees is typically based on the foraging distance of the bee community of interest. In the Mid-Atlantic USA, the mean foraging range of wild bees is approximately 500 m (Bartomeus et al. 2013; Kammerer et al. 2016), which, assuming equal foraging in all directions from a central nest, corresponds to a landscape of approximately 79 ha. In this region, the mean area of a habitat patch is only about 0.17 ha, or 0.2% of the 79-ha landscape (M Kammerer, *unpublished analyses*) illustrating that conservation practices implemented in one patch of habitat (site) improve a very small proportion of the landscape that pollinators experience. Empirical data are lacking, but based on common suggestions of adding habitat in roadsides, field edges, or other marginal land (Hopwood 2008; Morandin and Kremen 2013), it is likely that most pollinator plantings in the Mid-Atlantic USA are quite small (< 1 ha), and likely inadequate to solely support whole bee communities (McCullough et al. 2021).

If landscape quality is a primary driver of bee communities, and conservation practices improve only a small section of a landscape, how likely are conservation practices to benefit wild-bee populations? A quantitative comparison of local and landscape effects is necessary to decide not where, but if adding small patches of wild-bee habitat is likely to realize a measurable increase in bee abundance or richness (Gonthier et al. 2014), although it is challenging to evaluate on a larger scale how populations are affected (Kleijn et al. 2018; Scherber et al. 2019). Quantitative syntheses of agroecosystems across the globe found abundance of wild bees increased with more complex landscapes and diverse local plant communities or vegetation types (Kennedy et al. 2013; Shackelford et al. 2013). Richness of all wild bees also increased with landscape quality, and, for solitary bees only, locally diversified fields supported more species (Kennedy et al. 2013). Comparing the effect of different scales, some studies have shown that local context matters more than landscape quality (Coutinho et al. 2018; Rollin et al. 2019), but there are also examples where landscape effects dominated (Bartholomée et al. 2020; Griffin et al. 2021; Coutinho et al. 2021) or compensated for intensively managed agriculture or local contexts that are otherwise challenging for wild-bee communities (Rundlöf et al. 2008; Papanikolaou et al. 2017).

Between spring and summer, composition of wild-bee communities changes substantially, which could lead to seasonal variation in importance of local and landscape resources. In the mid-Atlantic USA, there is substantial turnover in bee species present in spring (before mid-May) versus summer and fall (Kammerer et al. 2021; Turley et al. 2022). With changes in composition of the bee community comes seasonal variation in several important functional traits including voltinism, overwintering location, and body size (Osorio-Canadas et al. 2018). Body size is strongly linked to species’ typical foraging distances (Greenleaf et al. 2007), likely dictating the relative importance of local and landscape resources. However, to our knowledge, no studies have examined seasonal variation in dependence of bees on local vs. landscape resources. Moreover, habitats vary widely in the magnitude and timing of floral and nesting resources that they provide for bees (Ogilvie and Forrest 2017), which can lead to temporally variable relationships between wild-bee communities and land use (Cole et al. 2017; Galpern et al. 2021)

To facilitate translating results of landscape-scale studies to applied management and conservation, pollination ecologists should quantify landscape quality based on specific, seasonal resources and risks. But due to the complexity of measuring flowers, nesting, or pesticide risk at the landscape-scale, these factors have been understudied (but see Guezen and Forrest 2021; Smart et al. 2021; Bloom et al. 2021). Instead, most studies have described landscape quality using broad landscape metrics, such as percent semi-natural habitat (Ricketts et al. 2008; Kennedy et al. 2013). Broad land-use metrics are correlated with many aspects of bee health and community composition (Kennedy et al. 2013; Martínez-Núñez et al. 2020; Du Clos et al. 2020), but are less useful for applied conservation decisions, as land use is an indirect, rather than direct, driver of bee abundance and diversity (Roulston & Goodell, 2011). Land-use patterns dictate floral and nesting availability (Williams and Kremen 2007), pesticide risk, and disturbance regimes (mowing, tillage, logging), so representing landscape quality based solely on amount of semi-natural habitat cannot untangle relative importance of multiple drivers. Furthermore, different semi-natural habitat types vary in how many food resources they provide (Bartual et al. 2019), and documenting bee responses to broad land-use patterns precludes determining which habitats are most important for bees and how and when to offset resource scarcity.

To address these knowledge gaps and inform conservation practices for wild bees, we studied season-specific relationships, over a two-year period, between local and landscape quality and wild-bee communities at 33 sites in the Finger Lakes region of New York, USA. We paired field surveys of wild bees, plants, and soil characteristics at each site with a multi-dimensional assessment of landscape quality, including landscape composition, configuration, insecticide risk, and data generated from a novel, spatio-temporal evaluation of floral resources at the local and landscape scales (described in Iverson et al, in review for this issue). As part of our local-scale assessment, we measured soil characteristics because many species of wild bees nest in the soil (Harmon-Threatt 2020) and soil fertility influences floral abundance, quality and quantity of floral rewards, and resulting bee visitation (Carvalheiro et al. 2021). We included insecticide risk to wild bees in our quantification of landscape quality as myriad evidence shows insecticide exposure can negatively influence bee behavior and reproduction and that insecticides used locally can have far-reaching consequences (Goulson et al. 2015; Long and Krupke 2016).

We ask the following research questions: 1) What is the relative importance of local versus landscape scale in driving bee richness and abundance? 2) Which metrics of local and landscape quality best explain wild-bee abundance and richness? And 3) How do the relationships in (1) and (2) vary with season? Within these broad objectives, we were also able to identify which types of semi-natural habitat were the most important for spring and summer bee communities and how temporal variation in floral resources at local and landscape scales affects abundance and richness of wild bees. Due to a higher prevalence of cavity-nesting, large-bodied social species with longer foraging ranges in summer bee communities, we predicted that summer bee abundance and richness would be strongly linked with landscape quality, while smaller-bodied spring bees would respond more strongly to local floral resources and soil characteristics. Drawing on other studies in our region documenting limited or no effect of insecticide risk on wild bees at a landscape scale (McNeil et al. 2020; Kammerer et al. 2021), we expected that land use and floral area metrics would be the best predictors of wild-bee communities.

## 2. Materials and methods

### 1. Study region and site selection

We studied wild-bee communities at 33 sites in the Finger Lakes region of New York, USA (Figure 1). The Finger Lakes region is in south-central New York, and is approximately 42% semi-natural land (almost all forest habitats), 8% developed, and 49% agriculture, including pastureland (USDA NASS 2018). There is a regional gradient of landscape composition, with high forest cover in the south, and increasing agricultural land moving north and closer to the lakes. (Figure 1). In proximity to Seneca and Cayuga lakes, the climate is relatively moderate, particularly well-suited for specialty crops like wine grapes and tree fruit. Generally, in this region, crop diversity is quite high, especially on small farms, which are common (USDA NASS 2019). Twelve percent and 42% of farms are smaller than 4 ha and 20 ha, respectively (USDA NASS 2019). The small farm sizes and relatively high amounts of semi-natural habitat means many agricultural areas in our study region could still be considered ‘complex’ landscapes (Tscharntke et al. 2005). For example, within 1 km, only five of our 33 study sites had less than 20% semi-natural habitat, and none were below 1% semi-natural habitat (a ‘cleared’ landscape as defined by Tscharntke et al. (2005).

**Figure 1:**
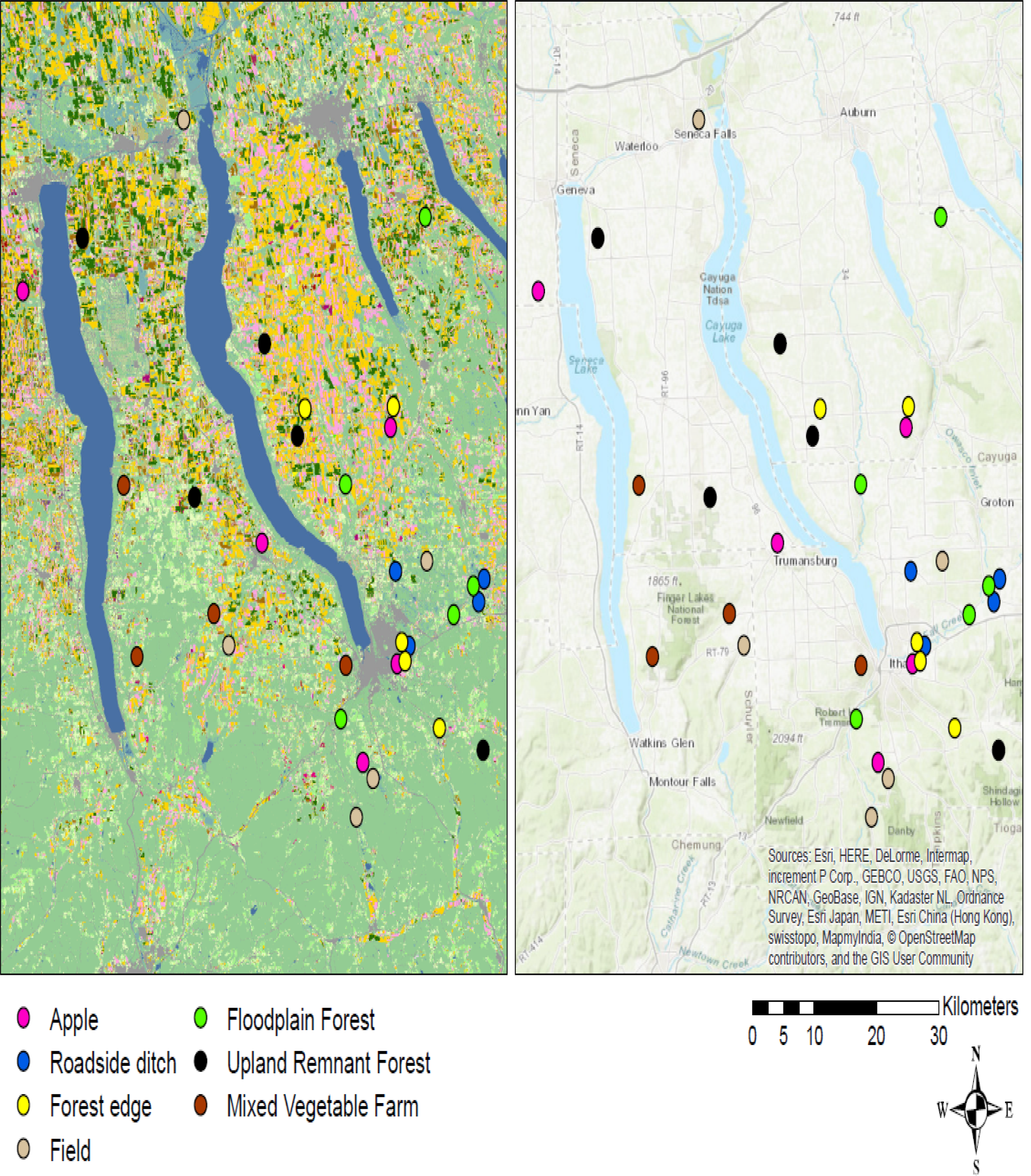
Map of study sites in the Finger Lakes region of New York, USA. Bees were sampled in spring and summer 2018-2019 at 33 sites in seven habitats defined by Iverson et al (in review; mesic upland remnant forests, floodplain forests, forest edges, old fields, roadside ditches, mixed vegetable farms, and apple orchards).

We selected our 33 study sites from 144 locations included in a previous study documenting plant community composition, local and landscape floral resources (Iverson et al.) in review, see Appendix for methodology). From these sites, we selected those corresponding to seven habitat types (mesic upland remnant forest (i.e., not previously cleared for crops), floodplain forest, forest edge, old field, roadside ditch, mixed vegetable farm, and apple orchard) that span a range of semi-natural to managed land use. Forest edge plots were placed adjacent to the boundary of upland forest and neighboring habitat (usually cropland), old fields were post-agricultural areas dominated by grasses and goldenrod (Solidago spp.; *Euthamia graminifolia*), and mixed vegetable farms were relatively small-scale farms that grew a diversity of fruits and vegetables. We selected the 33 sites from the initial 144 locations in Iverson (in review) based on attaining a sufficient sample size of each of the seven selected habitat types and on maintaining a minimum distance between sites. To ensure we were sampling independent bee communities, we chose sites that were, at minimum, one km from all other sites which, in our study region, exceeds the mean foraging range of a typical community of wild bees (Kammerer et al. 2016).

### 2. Wild-bee survey

At each of our 33 sites, in 2018 and 2019, we measured bee species richness and abundance using bee bowls according to the protocol of a long-term bee monitoring program in our region (Droege et al. 2016). In each year, we sampled bees in late April/early May and again in mid-July; dates were selected to correspond to peak floral abundance in forest, wetland, and successional habitats (Iverson et al. in review). We filled fluorescent blue, fluorescent yellow, and white, 355 mL Solo polystyrene plastic cups with 50:50 mix of propylene glycol and water. We placed bee bowls at the height of dominant vegetation for 7-14 d of sampling. In spring, we deployed bee bowls for 14 d, while, in summer, trap liquid evaporated more quickly, which limited our sampling to 7 d. At each site, we arranged bee bowls in 80-m transects in visible areas, alternating bowl color with 10 m between each bowl, for a total of 9 bowls per site. We placed the middle of our transects at the center of Iverson et al.’s (in review) plant survey plots, based on the recorded GPS coordinate, or, at some locations, physical flags marking the plot outline.

After collection, we stored bee specimens in 70% ethyl alcohol solution until pinning and sorting. We washed, pinned, and identified bee specimens, to species when possible, with taxonomic assistance from collaborators, including Dr. David Biddinger (Penn State Center for Pollinator Research, Biglerville, PA), and Dr. Rob Jean (Environmental Solutions & Innovations, Inc., Indianapolis, IN). We sorted specimens in the *Nomada* “bidentate” group to morphospecies, as this group is poorly resolved in existing taxonomic keys (Droege et al. 2010; Ascher and Pickering 2013). Also, some *Lasioglossum* specimens were damaged during collection or processing (n=50), so we could not reliably determine species identity. We excluded these specimens from richness analyses that required species-level identification. For all species, voucher specimens were deposited in the Frost Entomological Museum at The Pennsylvania State University in University Park, PA.

### 3. Local quality metrics

For each site, we measured soil characteristics and calculated plant species richness, community composition, and floral area from existing data (Iverson et al. in review).

#### 1. Soil collection and processing

In May 2018, we collected soil from each of our 33 study sites. Along the bee sampling transect, we collected five soil samples with a bucket auger to a depth of 9-18 cm, depending on rock and moisture content. We took shallower soil cores (9-12cm) at sites with very rocky or wet (floodplain habitat) subsoil. Wild-bee nesting would likely be inhibited by very high rock content or completely saturated subsoils (Harmon-Threatt 2020), so we considered the shallower sampling depth representative of the most favorable zone for bee nests. To quantify bulk density, at two locations along the bee transect, we collected three undisturbed soil cores (0-3 cm, 4-6 cm, and 7-9cm deep) with a slide hammer sampler (Soilmoisture Equipment Corp, Goleta, CA). Due to high moisture or rock content at some sites, we were only able to collect two bulk density cores, but we recorded the number of cores in each sample. For processing and analysis, we combined bulk density cores from all depths.

After collection, for all soil samples, we measured wet mass, then dried samples in an oven at 60°C for five days (or until the mass did not decrease) and measured dry mass. We calculated bulk density from dry mass and sent bucket-auger samples to the Penn State Agricultural Analytical Services Laboratory, where they measured pH, P, K, Mg, Ca, Zn, Cu, S, total nitrogen by combustion, percent organic matter, and percentage sand, silt, and clay via standard laboratory methods (Penn State Agricultural Analytical Services Lab). To summarize trends in soil characteristics, we centered and standardized soil variables and conducted a principal components analysis with the stats package in R (R Core Team 2021).

#### 2. Plant species richness, abundance, community composition, and floral area

We designed our study to leverage an existing, comprehensive plant survey conducted in the greater Ithaca region, NY (Iverson et al. in review, see Appendix). In 2016 and 2017, Iverson et al. (in review) documented plant species richness and abundance at 144 sites across the most common habitat types (N=22) in the surveyed region that span across the broader classes of forest, agriculture, wetland, successional, and developed. Briefly, Iverson et al. (in review) surveyed 3-20 sites of each habitat type based on variability among sites, with a median of ∼5 sites/habitat type. They used a Modified Whittaker plot design (Stohlgren et al. 1995) with halved dimensions, equating to a 10 × 25 m plot with nested subplots of varying sizes. They recorded plant cover by species in ten 0.25 m^2^ quadrats and species presence in all other subplots and in the full plot. Overall plant coverage was estimated based on frequency in larger subplots and coverage in the 0.25 m^2^ quadrats. Furthermore, they recorded the species abundance of all mature, i.e., potentially flowering, angiosperm trees.

Using a combination of survey data and flower density and size measurements for each species observed, Iverson et al. (in review) estimated floral area per species in each site. To estimate flower density, for each species, they measured the number of flowers in a 0.25 m^2^ quadrat during peak bloom and measured flower size in the field, from herbaria specimens, or from published sizes in online plant databases. Then, utilizing bloom dates published in a flora specific to the region, they estimated FA across the bloom window of each plant species. Finally, they calculated the daily average FA per m^2^ of each of the sampled habitat types by averaging the sum of the FA, weighted by percent cover, of each of the flowering plant species present across all plots in each habitat type (excluding apple blossoms in ‘apple orchard’ habitat, in order to capture only orchard-floor floral resources).

For our analysis, we summarized site-level FA curves with seven metrics: season total FA, minimum FA, maximum FA, FA coefficient of variation, and total FA in spring, summer, and fall. For total, maximum, and coefficient of variation, we summarized FA from April to mid-November. We quantified minimum FA from a narrower timeframe associated with more rapid plant growth (mid-May to mid-September) to avoid all sites receiving similarly low values associated with the beginning or end of the season. We defined ‘spring’ as early April to mid-June (day-of-year 92 to 163), ‘summer’ as mid-June to late August (day-of-year 164 to 238) and ‘fall’ as late August to mid-November (day-of-year 239 to 310). We generated two versions of each FA metric, one that represents the FA of all plants, and the second that only includes plant species that are known to be pollinated by insects (‘IP plants’, Iverson et al. in review).

From the plant survey data, we also quantified plant species richness, relative abundance, and community composition for our 33 study sites. We calculated species richness of all plants, species richness of insect-pollinated plants, and percent cover of all vegetation. We used non-metric multi-dimensional scaling (NMDS) ordination with a Bray-Curtis distance measure to quantify the main gradients in plant-community composition. Specifically, with the *vegan* package in R (Oksanen et al. 2013), we summarized the similarity between plant communities at all 144 sites surveyed by Iverson et al. (in review) and visualized the results in two dimensions. To determine the effect of varying composition of plant communities on bee abundance and richness at our 33 study sites, we included scores from the first two NMDS axes as predictor variables in our analyses.

### 4. Landscape quality metrics

#### 1. Landscape composition and configuration

To describe the land cover surrounding our study sites, we used a high-resolution map of land cover available for our study region (Iverson et al. in review). This product utilized a regional 1m resolution dataset of land cover (Chesapeake Conservancy 2013) to differentiate impervious surface, trees, and low vegetation and a regional natural habitat map (Ferree and Anderson 2013) and the USDA Cropland Data Layer (USDA NASS 2018) to resolve more detailed natural and agricultural habitats, respectively. Wetland habitats were incorporated using the National Wetlands Inventory data (U.S. Fish and Wildlife Service 2017) and a ‘roadside ditch’ habitat class was created by adding a 3m buffer on either side of non-urban roads.

From this high-resolution land-cover map, we calculated landscape composition and configuration within 1 km radius of each of our sampling sites. We selected this distance because, excluding large-bodied *Bombus* sp., most wild-bee species in our region forage within 1km of their nests (Kammerer et al., 2016). We grouped land-cover classes to form six metrics of landscape composition: percentage of the landscape in agriculture, forest, successional (old field and shrubland), wetlands, water, and developed habitats (Table S1). Based on previous research examining landscape configuration effects on wild bees (Kennedy et al. 2013), we also calculated six landscape configuration metrics to represent the aggregation, shape, and diversity of habitat patches around our sampling sites. Specifically, using the *landscapemetrics* package, version 1.4.2, in R (Hesselbarth et al. 2019; R Core Team 2021), we calculated edge density, Shannon diversity of land cover classes, Simpson diversity of land cover classes, mean perimeter-area ratio, variation in nearest neighbor distance between patches of the same class, and interspersion and juxtaposition indices.

#### 2. Landscape insecticide toxic load

To represent risk to wild bees from insecticide applied to agricultural crops, we calculated an index of insecticide toxic load. We generated the index of insecticide toxic load (Douglas et al. 2021) using our high-resolution land cover maps and insecticide data from 2014, as this is the most recent year with a complete, publicly available dataset of insecticide application. We estimated insecticide risk at our sampling sites in three steps. First, we calculated a distance-weighted metric of landscape composition from the high-resolution land cover data (see *Landscape composition and configuration*) with the ‘distweight_lulc’ function in the *beecoSp* R package (Kammerer and Douglas 2021). For distance-weighting, we selected a wild-bee foraging range of 500 m, which assumes, from the center of each study site, 70% and 100% of bee foraging occurs within 500-m and 1-km radii, respectively. Then, for each land-cover class, we multiplied distance-weighted area by the insecticide-toxic-load coefficient for a given land cover. Finally, for each study site, we calculated total insecticide toxic load as the sum of insecticide load from all land cover classes within a 1-km landscape. Without *a priori* knowledge of the most likely route of insecticide exposure for each active ingredient, we used insecticide values corresponding to the mean of oral and contact toxicity (Douglas et al. 2021).

#### 3. Floral area of landscapes

We quantified the floral area of landscapes in three steps. First, we averaged floral area at all sites within a land-cover class to calculate habitat-level FA per day per ha of habitat. Then, for each land-cover class within 1km of our study sites, we multiplied distance-weighted area (ha) in the landscape (see *Insecticide toxic load* for distance-weighting details) by the habitat-level FA. Then, we summed over all land-cover classes, yielding landscape-total FA per day for each of our study sites. For local and landscape floral area, we calculated minimum FA, maximum FA, FA coefficient of variation, and total FA in spring, summer, and fall for all plants and for insect-pollinated plant species only (‘IP plants’, Iverson et al. in review).

#### 4. Distance to water and topography

We generated distance to water and topography metrics and included these factors in our analyses. To differentiate riparian sites very close to water, we calculated the minimum distance from our sampling locations to water (streams, rivers, ponds, or lakes). We identified water features closest to our sampling sites using the National Hydrography Dataset (U.S. Geological Survey 2019a) for New York state. Lastly, we included topographic information to represent the micro-climate at each of our sites. We calculated elevation, slope, and aspect at each sampling site from the 1/3 arc-second (approximately 10m) resolution USGS National Elevation Dataset (U.S. Geological Survey 2019b). We included distance to water and topography as landscape variables because, excepting elevation, they describe the relationship between the sampling site and surrounding water and topographic features. However, distance to water and topography were not among the most explanatory predictor variables (Figure 2), so if we had classified them as local variables, our results would be largely unchanged. We used the R statistical and computing language (R Core Team 2021), specifically the *sf* and *raster* packages (Pebesma 2018; Hijmans 2022), for all manipulations and analyses of geospatial data.

**Figure 2:**
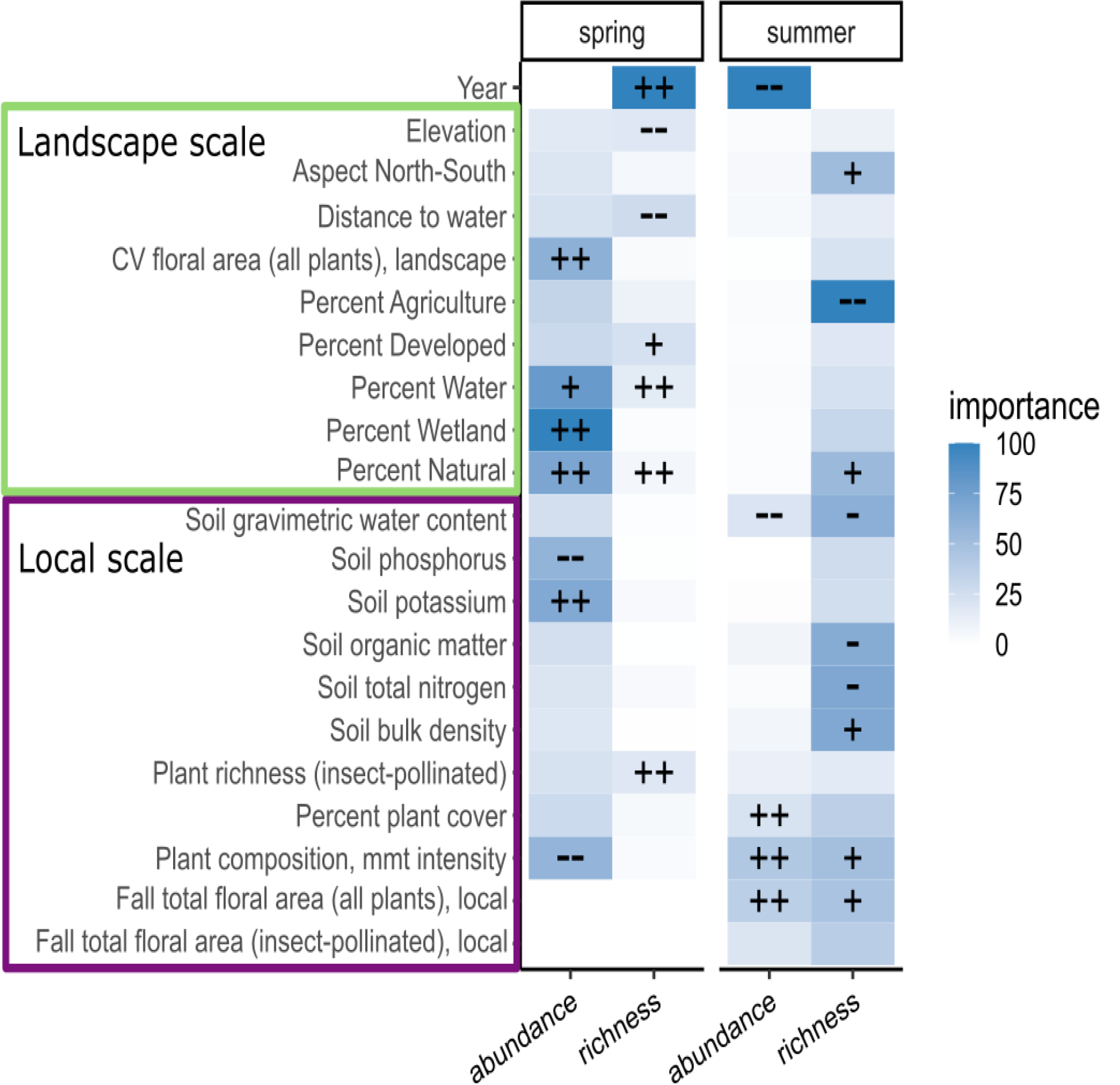
Variable-importance scores of the top 21 variables from random forest models predicting wild-bee abundance and richness. Bees were sampled in 2018 and 2019 at 33 sites in the Finger Lakes region of New York, USA. We calculated landscape variables for area within 1km of our study sites. For the most important variables, symbols denote directionality of the relationship (‘++’ = moderate/highly positive, ‘+’ = slightly positive, ‘-’ = slightly negative, and ‘-–’ = moderate/highly negative). Variable abbreviations are as follows: ‘CV floral area’ = coefficient of variation of floral area, ‘Plant composition, mmt intensity’ = composition of plant community associated with a gradient in management intensity (Figure S1).

### 5. Statistical analyses

#### 1. Sampling-effort adjustment

Prior to analysis, we adjusted bee abundance and species-richness measures to account for varying sampling effort. While we were collecting bee bowls, we recorded the number of traps that were cracked, tipped over, or otherwise compromised. We present abundance results as bees per successful trap per day, to adjust for varying sampling time and lower sampling effort at sites where traps cracked or fell over. Unfortunately, for the July 2019 sampling round, we lost our record of the number of compromised traps. To the best of our ability, we recreated these data from memory shortly after the sampling period by estimating the number of compromised traps per site. For wild-bee richness analyses, we adjusted for uneven effort using coverage-based rarefaction and extrapolation (Chao and Jost 2012), rather than the number of successful traps, so recreated data were not used in richness analyses. Specifically, we estimated bee richness at the mean coverage level for each season using the *iNEXT* package in R (Hsieh et al. 2016, 2018; R Core Team 2021).

#### 2. Random forest models

We used random forest models to compare the relative importance of local vs. landscape variables, identify the most important individual predictor variables, and examine relationships between predictors and wild bee abundance and richness. Random forest models are robust to correlated predictors and can represent complex, non-linear relationships, which we expected in our dataset (Wright and Ziegler 2017). In our study region, the composition of wild-bee communities in spring is substantially different from summer and fall (Kammerer et al. 2021; Turley et al. 2022), so we analyzed our spring and summer sampling periods separately. For spring analyses, we removed floral-area metrics that represent, or were correlated with, floral resources available in summer and fall (Figure S10). Specifically, for spring analyses, we removed summer-total-floral area and fall-total-floral area of all plants and insect-pollinated (‘IP’) plants, maximum-floral area (IP plants) and total-floral area (IP plants).

In preliminary analyses, we noticed that the number of predictor variables included in random forest models influenced model performance and variable importance scores (Fox et al. 2017). To compare random forest models with the same number of predictors, we first modelled bee abundance and richness using all 72 year, landscape, and local predictors and examined the resulting variable importance scores. Then, we selected the top 35 predictors and generated a final model using only these predictors. We used ‘top-35’ models to compare variable importance and describe relationships between bee abundance, richness, and specific predictors (Figures 2-5). We assessed the performance of all random forest models using 10-fold cross validation repeated 15 times, and, with a grid search, selected the optimum number of trees (1000 to 5000, incremented by 1000) and variables at each tree split (15 to 30, incremented by 5). We used the random forest algorithm from the *ranger* package in R (Wright and Ziegler 2017), tuned the model with the *caret* package (Kuhn 2008, 2019), and, to examine our results (Figures 3-5), generated accumulated local effects (ALE) plots with the *iml* package (Molnar et al. 2018). ALE plots depict the relative effect of changing one predictor variable and are centered at zero so each value on the curve is the difference from the mean prediction of the random forest model. For each season, we present local effect plots for year (Figure 5) and the top four variables predicting bee abundance and richness (Figures 3-4).

**Figure 3:**
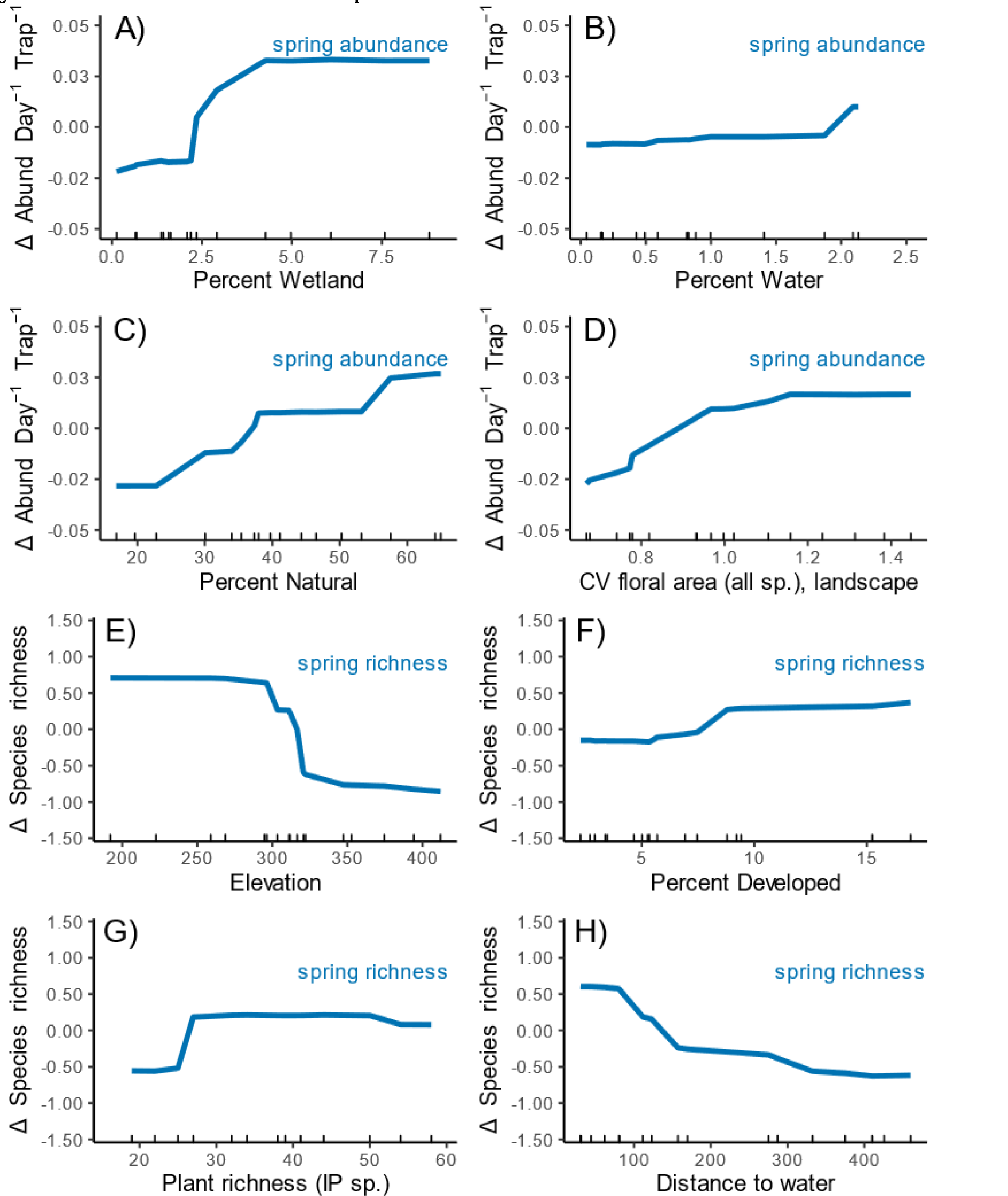
Relationship between landscape and local predictors and wild-bee abundance and richness in the spring. For both abundance and richness models, we show the top four predictors (excluding year, shown in Figure 5), with x-axis truncated to 10-90% quantiles (n=29 sites). ‘IP’ indicates insect-pollinated plants. To enable comparing relative effects across seasons with varying mean abundance or richness (Table 1), we depict values on the y-axis as difference from the predicted mean.

**Figure 4:**
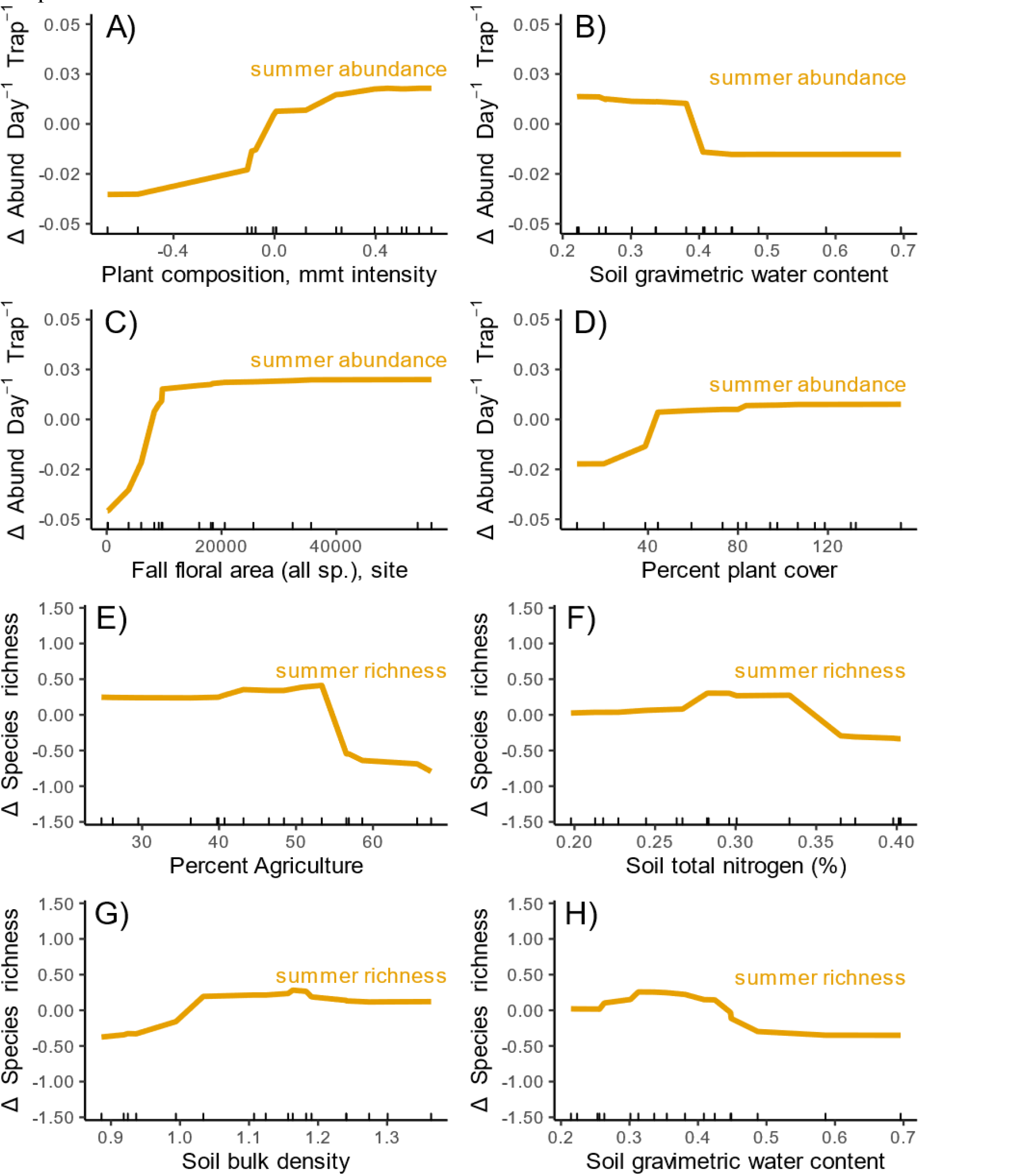
Relationship between landscape and local predictors and wild-bee abundance and richness in the summer. For both abundance and richness models, we show the top four predictors (excluding year, shown in Figure 5), with x-axis truncated to 10-90% quantiles (n=29 sites). To enable comparing relative effects across seasons with varying mean abundance or richness (Table 1), we depict values on the y-axis as difference from the predicted mean.

**Figure 5:**
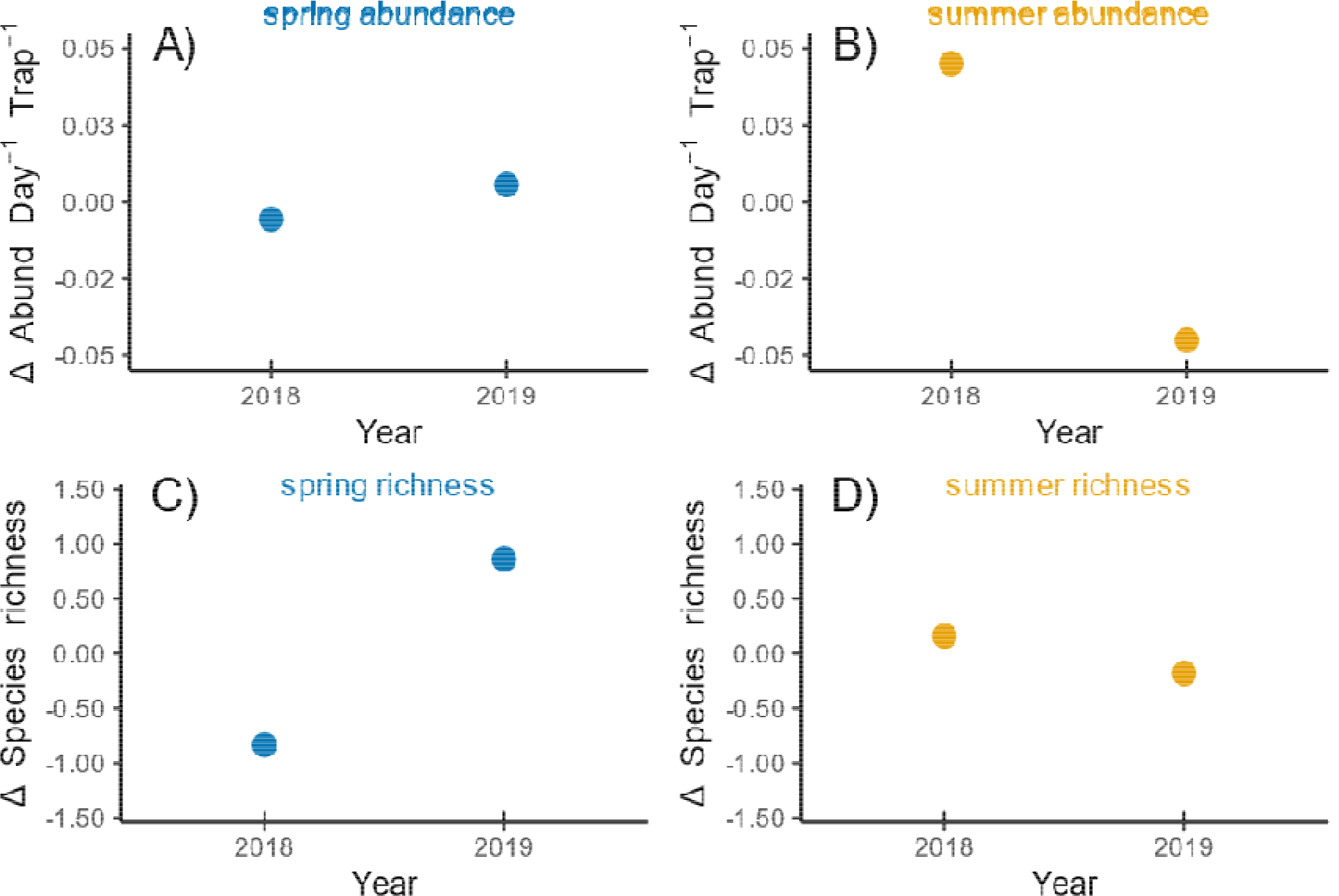
Difference between 2018 and 2019 for abundance (top) and richness (bottom) of wild bees in spring and summer estimated using a random forest model. To enable comparing relative effects across seasons with varying mean abundance or richness (Table 1), we depict values on the y-axis as difference from the predicted mean.

## 3. Results

### 1. Wild-bee communities

In spring and summer, we documented diverse wild-bee communities. Our survey yielded 3108 specimens, 1666 in 2018 and 1442 in 2019. We collected more bees in spring (n=2130) compared with summer (n=987), particularly in the second year when our summer sampling generated only 282 individuals. We documented 127 species of wild bees of 21 genera, with 94 species present in spring and 78 in summer. *Andrena, Lasioglossum*, and *Ceratina* were the most abundant spring genera, representing 78% of the individuals we observed (Figure 6). At the species-level, the most abundant spring bees (> 5% spring abundance) were *Andrena carlini* Cockerell, *Ceratina calcarata* Robertson, *Osmia cornifrons* Radoszkowski, *Ceratina dupla* Say, and *Andrena hippotes* Robertson. In summer, *Lasioglossum* was the most abundant genus, followed by several other genera in the family Halictidae (*Agapostemon, Augochlora, and Halictus*). *Lasioglossum leucozonium* Schrank, *Agapostemon virescens* Fabricius, *Lasioglossum versatum* Robertson, *Peponapis pruinosa* Say, and *Augochlora pura* Say were the most abundant summer species (> 5% of summer abundance).

**Figure 6:**
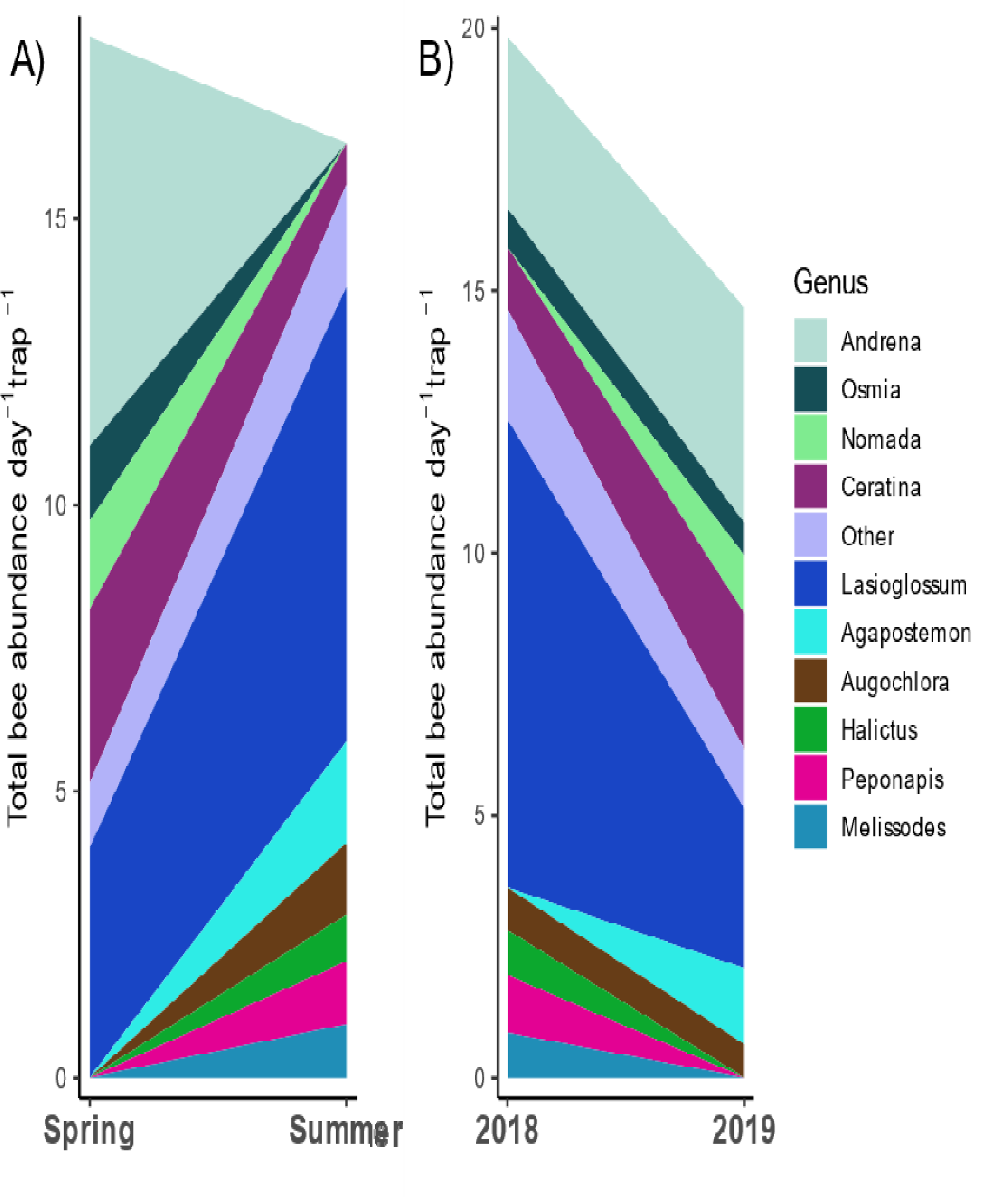
Abundance of wild-bee genera by season (panel A) and year (panel B) at 33 sites in the Finger Lakes region of New York, USA.

### 2. Local quality

#### 1. Characterizing local quality

We characterized several dimensions of local quality for wild bees, including floral area, plant richness and community composition, and soil characteristics. Of the seven habitats included in our study (mesic upland remnant forest, floodplain forest, forest edge, old field, roadside ditch, mixed vegetable farm, and apple orchard), mesic upland remnant forest and forest edge had highest floral area, peaking in early spring (Figure 7). In summer and fall, floral resources in all habitats were generally much lower, excepting an early-fall peak in flowers in old fields associated with goldenrod (*Solidago* spp. and *Euthamia graminifolia*) bloom period (Figure 7, Figure S3).

**Figure 7:**
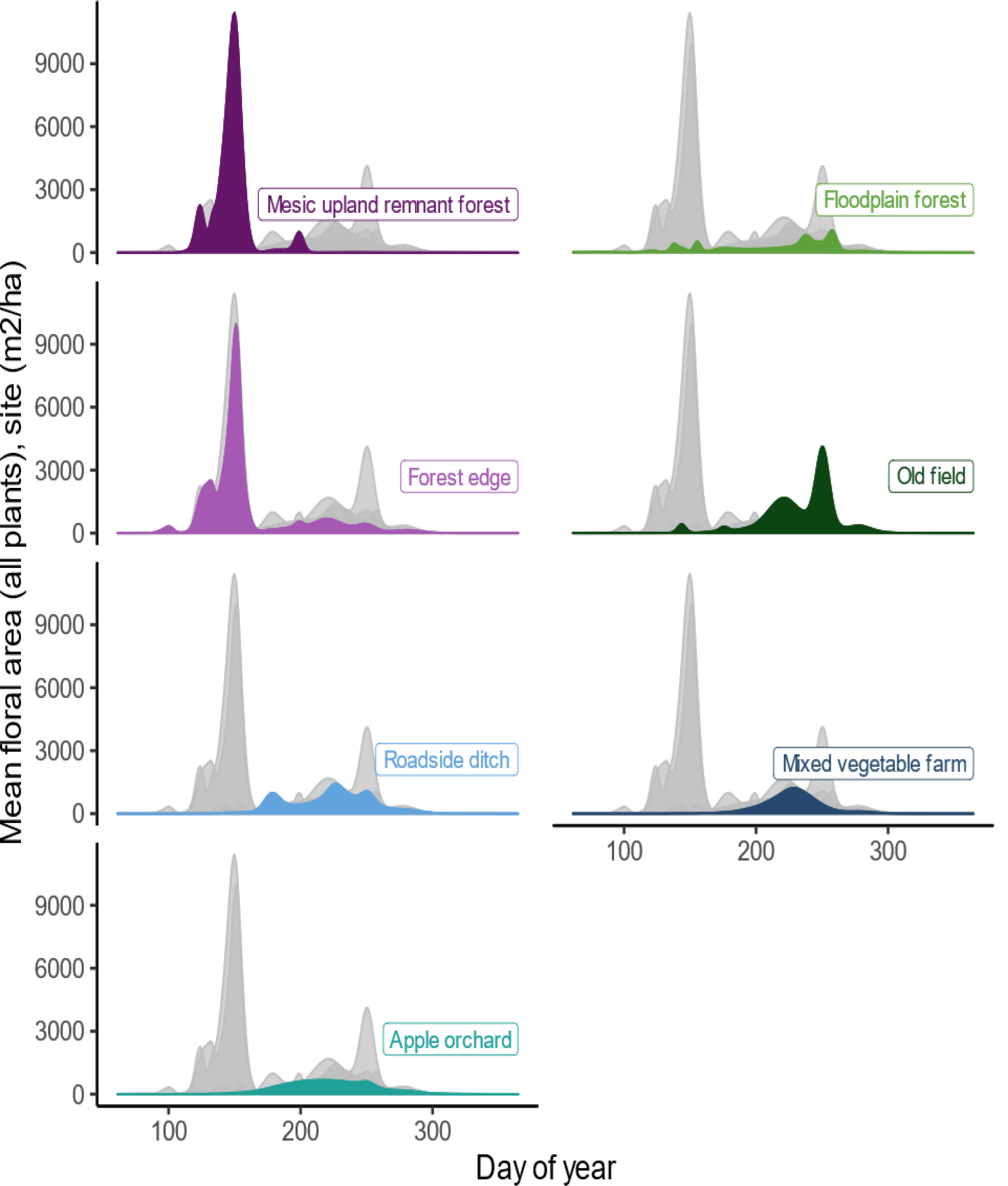
Floral area over time available in seven habitats in the Finger Lakes region of New York, USA. Floral area values represent all flowering plants. Floral area of the labelled habitat is represented with a colored polygon, while all other habitats are indicated with the grey polygons.

Our analysis of plant community composition revealed clear differentiation between plant communities in different habitats. We analyzed the full set of habitat types (n=22, Iverson et al. (in review) to capture broad gradients in composition of plant communities that were not evident when including only our focal habitats (n =7, M Kammerer, *unpublished analyses*). Considering all 22 habitats, the first ordination axis (‘NMDS1’) differentiated habitat types according to the intensity of human management (Figure S1, stress = 0.153), with unmanaged forest and wetland habitat separating from managed agricultural and developed habitats (lawn, row crops, orchards, vegetable farms, pasture, and hay fields). Old fields, roadside ditches, tree rows, and forest edge habitats were intermediate, containing plants associated with managed and unmanaged habitats. In subsequent random forest models, we included site values on NMDS axis one as a proxy for intensity of management. The second axis was less clear, but seemed to represent a gradient of moisture content, separating wetlands from forest and managed habitats, with floodplain forests in the middle (Figure S1).

We found significant variation in soil characteristics between the seven habitat types we sampled. There were two main gradients in our soil dataset revealed by the principal components analysis (Figure S2). Explaining 26.4% of the variation in our soil data, the first principal component was associated with pH, soil texture (percent sand, silt, and clay), water content, organic matter, and total nitrogen. Roadside ditches had more basic, sandier soil than any of the other habitat types, except some floodplain forest samples. All other habitat types had loam to silt-loam soil. Interestingly, soil texture was highly variable between floodplain forests, with soil texture in this habitat encompassing the full range of texture classes represented at all other sites. The second principal component explained 20.0% of variation in soil characteristics and correlated with soil clay content, potassium, copper, phosphorus, zinc, clay content, and cation exchange capacity. Specifically, some vegetable farm and orchard samples had much higher potassium, phosphorus, copper, and zinc content than the other habitats, likely due to fertilizer or manure application to support crop growth. Soils at forested sites were characterized by higher organic matter and total nitrogen, possibly from leaf litter accumulation.

#### 2. Local quality effects on wild bees

Comparing spring to summer, we documented substantially different relationships between local quality and bee abundance (Figure 2). In spring, we observed moderate effects of local characteristics on wild bees, with three local variables (soil potassium, soil phosphorus, and management intensity as reflected in plant community composition) among the top predictors of abundance or richness. Abundance of spring bees increased with higher levels of soil potassium and lower soil phosphorus measured at the site, with the greatest increase between approximately 75 and 125 ppm potassium (Figure S9 B-C). Abundance of spring bees was lower in more-managed sites (Figure S9 A). In summer, abundance of wild bees was influenced by the level of management (plant community composition) and presence of fall-flowering plants (Figure 2). At the less-managed sites (remnant forests and floodplain forests, NMDS axis one loading less than approximately - 0.1), we observed lower abundance of summer bees (Figure 4A). Bee abundance was intermediate in forest edge, old field, and ditch habitats (NMDS1 =-0.1 to 0.4), and highest in orchards and mixed vegetable farms (NMDS1 > 0.4). Sites with very low or no fall-flowering plants also had notably lower abundance of summer bees, but, when floral area was greater than approximately 10,000 m^2^/ha of habitat, bee abundance did not increase with additional fall flowers (Figure 4C).

For richness of wild bees, local quality had weak effects on spring and summer bees (Figure 2). In spring, we observed approximately one additional bee species at sites with more than 25 plant species. In summer, soil fertility, bulk density, and water content were the most important local predictors. We documented slightly more species of bees at sites with intermediate total soil nitrogen, bulk density, and water content, peaking at approximately 30%, 1.18 g/cm^3^, and 0.31 g/g, respectively (Figure 4F-H).

### 3. Landscape quality

#### 1. Characterizing landscape quality

We noted substantial variation in the composition and configuration of landscapes surrounding our study sites (1 km radius), while topography was more consistent (Figures S5-S7). In our study landscapes, amount of developed land ranged from zero to approximately 50%, while forest and natural habitats ranged from zero to more than 70%. All landscapes surrounding our study sites were at least 18% agriculture, up to a maximum of 91% agricultural land. Distance to the nearest water source was generally low, with a maximum of 580 m. Most of our study sites had relatively low insecticide toxic load values, but insecticide toxicity was substantially higher in landscapes with significant apple and grape production (Figure S5). Edge density was the most variable configuration metric, while the interspersion and juxtaposition index and perimeter-area ratio were more consistent. Both Shannon and Simpson diversity indices were left-skewed, capturing the diverse agricultural, natural, and developed land cover types present in our study region (Figure S6).

#### 2. Landscape quality effects on wild bees

In spring, we documented higher abundance of wild bees at locations with more wetland, surface water, natural habitat, and flowers in the surrounding landscape (Figure 3A-D). In our study region, wetlands and surface water are a relatively small percentage of most landscapes (3.42% ± 3.54%, mean ± standard deviation of wetland area), but we found substantially higher abundance of spring bees at sites with 4-5% wetland in the landscape compared with those that had 0-2% wetland (Figure 3A). Between approximately 20 and 60% natural land in the landscape, abundance of spring bees increased, likely due to additional floral resources and nesting sites (Figure 3C). Lastly, we observed higher abundance of spring bees with more variable (higher CV) landscape-level floral area (FA, Figure 3D). In our study region, high variability in FA comes from multiple, high peaks in spring FA from flowering trees in forested habitats (Figure 7).

For species richness of spring bees, the amount of developed land, elevation, and distance to water in the landscape were among the best predictors. We found more species at low elevation sites and locations with some developed land in the landscape, although most of our study landscapes had a relatively small amount of developed land (less than 10%) (Figure 3E-F). At sites less than 100 m from water, we observed approximately two more species of spring bees than sites that were 400 m or more from the nearest stream, lake, river, or pond (Figure 3H).

Compared with spring, in summer, landscape effects on wild bees were moderated. We did not detect any relationship between landscape quality and abundance of summer bees, but three landscape variables (percent agriculture, percent natural, and north-south aspect) were among the most important predictors of richness of summer bees (Figure 2). We observed a threshold of approximately 55% of the landscape in agriculture, above which richness of summer bees dropped substantially (Figure 4E). Insecticide toxic load, slope, and landscape configuration metrics were not among the most important predictors of spring or summer bee communities.

#### 4. Relative importance of year, local, and landscape quality for wild bees

Generally, with our 72 local and landscape-quality metrics, we were able to explain a substantial amount of the variation in wild-bee abundance and richness. Modeling abundance of bees in both seasons, we explained more than 38% of the variation in our data using year, local, and landscape variables (Table 1). We had more unexplained variation for species richness, with mean r-squared values of 20% and 28% for the best random forest models in spring and summer, respectively (Table 1). Across all models, however, our prediction error was very high (72%-108% of mean abundance or richness, Table 1), indicating we are unable to reliably predict bee communities in additional locations or years. As a result, we focused on presenting relationships among landscape and local quality and bee abundance and richness, rather than prediction.

**Table 1:** Performance of random forest models predicting wild-bee abundance and richness at 33 sites in the Finger Lakes region of New York, USA. We show mean abundance or richness, root mean-squared error (RMSE), r-squared (variance explained), and mean absolute error (MAE) as mean ± standard deviation from 10-fold cross validation, repeated 15 times. We tuned the number of variables at each tree split (mtry) and the number of trees (ntrees) with a grid search and present optimal values for each random forest model.

| Season | Response Variable | Mean Abundance or Richness | Variables Included in RF Model | RMSE | R-squared | MAE | mtry | ntrees |
| --- | --- | --- | --- | --- | --- | --- | --- | --- |
| spring | Abundance Day<sup>-1</sup> Trap<sup>-1</sup> | 0.28 ± 0.27 | Landscape, Year, Site | 0.20 ± 0.10 | 0.46 ± 0.32 | 0.15 ± 0.06 | 25 | 5000 |
| | Richness | 11.79 ± 8.60 | Landscape, Year, Site | 8.07 ± 3.33 | 0.20 ± 0.23 | 5.97 ± 2.20 | 25 | 5000 |
| summer | Abundance Day<sup>-1</sup> Trap<sup>-1</sup> | 0.25 ± 0.27 | Landscape, Year, Site | 0.21 ± 0.11 | 0.38 ± 0.24 | 0.16 ± 0.06 | 30 | 1000 |
| | Richness | 7.13 ± 7.24 | Landscape, Year, Site | 6.09 ± 2.76 | 0.28 ± 0.25 | 4.74 ± 1.76 | 15 | 5000 |

Comparing landscape, year, and local predictors, we found generally that landscape variables and year explained more of the variation in bee abundance and richness than local quality. However, the importance of local versus landscape scale was different for abundance and richness models and varied by season (Figure 2). For abundance of spring bees and richness of summer bees, landscape variables were the most important predictors (Figure 2). In spring, year was, by far, the most important predictor of species richness, with more species of spring bees in 2019 than 2018 (Figure 5). In addition to year, for richness of spring bees, we observed weak effects of landscape composition, but only one local variable was among the best predictors (Figure 2). Year was also the most important predictor of abundance of summer bees, but, unlike spring richness, local quality was almost as important (Figure 2).

## 4. Discussion

In this study, we found that bee abundance and richness responded to an interplay of landscape, local quality, and year with significant differences between spring and summer bee communities. Though we predicted spring bees would be more dependent on local quality than summer bees, we instead found that spring abundance and spring and summer richness were mostly associated with landscape quality, specifically landscape floral resources in the springtime and percentage of landscapes occupied by wetlands, water, natural, and agricultural habitats. Summer abundance was significantly affected by local conditions, including soil fertility, plant-community composition, and local-scale floral resources in the fall. Consistent with our previous studies, predicted insecticide risk in the landscape was not a significant factor for the wild bee communities in our study. Here, we discuss wild-bee responses to landscape, then local quality, with recommendations for designing conservation practices and suggestions for future research.

Wetlands and surface water were very important landscape features for supporting spring bees; therefore, we expect that restoring or conserving relatively small patches of these habitats could substantially improve landscape quality in spring. We observed more species of spring bees closer to water, indicating that terrestrial-aquatic interfaces likely support specialist bees or plant species important for spring bees (Stewart et al. 2017). This pattern underscores the short foraging distance of many wild-bee species in our study region, as bee richness decreased substantially only 200m removed from water. In our study region, wetland habitats, especially shrub and emergent wetlands, host a unique community of plants, including some herbaceous flowering plants not found in other habitats (Figure S1). Several of the plant species associated with floodplain and emergent wetland habitats are recommended for pollinator plantings (e.g. *Eutrochium purpureum, Helianthus decapetalus;* Byers et al., n.d.), or are closely related to known pollinator-attractive species (e.g. *Bidens cernua, Hydrophyllum canadense, Stachys hispida)*, suggesting they might also provide pollen or nectar resources for wild bees (Tuell et al. 2008). There are relatively few studies documenting wild-bee communities in wetlands, although most report rich bee communities in and around water and wetland habitats (Stewart et al. 2017; Vickruck et al. 2019), including presence of some threatened species (Moroń et al. 2008). In our study, relatively small areas of wetlands in a landscape correlated with higher bee abundance and richness, so we conclude that surface water and wetland habitats would be feasible targets of conservation programs seeking to improve landscape quality for wild bees. Also, wetlands host many other insect, birds, and amphibian taxa (Riffell et al. 2003; Gibbons et al. 2006; O’Neill and O’Neill 2010), and thus wetland restoration and conservation could contribute more broadly to biodiversity goals. These findings also confirm the importance of existing programs that protect wetland habitat from agricultural and urban development.

In spring, wild bees are heavily dependent on resources provided by the surrounding landscape, most likely from mass flowering of trees in forest habitats. We found more spring bees at sites with multiple, high peaks in spring floral resources from flowering trees in forests. Past research has documented a diverse wild-bee community in tree canopies (Urban-Mead et al. 2021) and demonstrated that solitary bees consume spring pollen from wind-pollinated trees (Kraemer and Favi 2010). In our study, spring bee abundance was positively associated with floral resources provided by the entire plant community (wind and insect pollinated plants) reinforcing the finding that these bees use trees not traditionally thought to be ‘insect pollinated’, such as oaks (*Quercus* spp.), hickories (*Carya* spp.), birches (*Betula* spp.), American beech (*Fagus grandifolia*), and maples (*Acer* spp.). The pattern we observed suggests that spring bees were able to take advantage of very large pulses of floral resources such that short periods of mass flowering may be equally beneficial as consistently high levels of flowering (Hemberger et al. 2020).

In summer, diversified vegetable farms and apple orchards hosted high abundance of bees, but not all species persisted in highly managed landscapes. In mid-summer, old fields had more floral resources than agricultural habitats (Figure 7), but plant communities in orchards and mixed vegetable farms hosted higher bee abundance. We speculate that, in these agricultural habitats, flowers of crop plants and weeds in fencerows, fallow, or between-row sections of fields are particularly important floral resources for bees. Indeed, many studies have documented the importance of weeds in providing forage for bees in both agricultural and urban areas (Norris and Kogan 2005; Bretagnolle and Gaba 2015; Requier et al. 2015; Rollin et al. 2016). Also, most of the farms and orchards we sampled were not intensively tilled or have substantial untilled area in proximity, which likely benefits species of ground-nesting bees (Ullmann et al. 2016). While some agricultural habitats can support relatively high bee abundance, like many other studies (Kennedy et al. 2013; Grab et al. 2018), we documented fewer species of summer bees in landscapes with a high proportion of agricultural land. In contrast to our sampled vegetable and orchard crop farms, most of the agricultural land in our study was intensively managed row crop agriculture. Similar to simplified bee communities in urban areas, crops and weedy plants in intensively managed agricultural habitats host a subset of all bee species, likely generalist, disturbance-tolerant species (Harrison et al. 2018).

We found that soil fertility was an important component of local habitat quality for wild bees, potentially due to fertility effects on floral resources. Intermediate soil fertility was associated with higher richness of wild bees in summer. We posit that soil fertility primarily affected wild-bee communities at our study sites by mediating plant vigor and flowering. Soil-nutrient levels influence richness and composition of plant communities (Tilman 1987; DiTommaso and Aarssen 1989; Wilson and Tilman 1991), but, we found little evidence that plant community shifts drove fertility effects on bees because plant richness was a poor predictor of bee abundance and richness, and plant community composition and soil fertility were weakly correlated (Pearson’s *r* = -0.6, -0.52, and 0.4 between NMDS axis 1 and soil organic matter, total nitrogen, and potassium, respectively). Other studies have shown that soil-nutrient levels influence size and number of flowers and quality and quantity of nectar and pollen produced, with measurable effects on bee visitation (Burkle & Irwin, 2010; Burkle & Irwin, 2009; Cardoza et al., 2012). Based on our results, manipulating soil nutrients within restoration and conservation plantings may increase floral resources and associated bee communities, but we need targeted experimental research to verify that patterns we documented are consistent and strong enough to be a successful management strategy without accompanying unintended consequences.

For both seasons, soil texture was notably absent from our list of important local variables. If, at the local level, wild-bee communities were limited by the availability of nests, we expected that soil texture would have been an important soil characteristic, as many wild bees nest in the ground or use soil for cavity partitions (Pinilla-Gallego et al. 2018; Harmon-Threatt 2020). In our study, soil texture within each study site was not a strong predictor of bee abundance or richness, suggesting that numbers of suitable nest sites may be governed by soil characteristics within the larger landscape (Hellwig et al. 2022), which we could not capture. Few studies have quantified landscape-scale nesting resources (Ariza et al. 2022), likely in part due to poor characterization of preferred nesting conditions of wild bees (Harmon-Threatt 2020) and challenges associated with existing soil maps (Cambardella et al. 1994; Moral et al. 2010). In the future, it would be extremely valuable to quantify, at the landscape-scale, availability of soil, wood, stem, and cavity nest sites for wild bees based on empirical data, mirroring the Iverson et al. (in review) effort to estimate landscape floral resources.

We observed notable differences between the two years of our study, with more species of spring bees and fewer summer bees in 2019 compared with 2018. In 2019, early spring was warmer than 2018 and we delayed our sampling time due to logistical constraints which likely altered the composition of the bee community we sampled. We attribute the higher richness of spring bees in 2019 to a better match between our sampling time and flight period of most spring bees (Kammerer et al. 2021). The differences in the summer collections was likely also due to weather conditions in the previous year, as the summer of 2018 was extremely rainy, with near-record high precipitation (DiLiberto 2018), which may have reduced foraging opportunities and negatively affected bee populations (Tuell and Isaacs 2010; Vitale et al. 2020). Evaluating the effect of weather conditions on bee communities requires multiple years of sampling, and thus we did not formally include these variables in this study, but such analyses have been conducted in other studies (Kammerer et al. 2021; Filazzola et al. 2021).

Though we were able to provide important insights about how local and landscape variable influence bee communities, our study has several limitations. First, we sampled wild bees using pan traps, which perform poorly in areas with abundant floral resources and have some taxonomic biases, including under-sampling large bodied bees (Roulston et al. 2007; Baum and Wallen 2011). By using pan traps, we were not able to determine if the bees we sampled were nesting or foraging within our study site, or en route to a different patch. We minimized, but could not completely exclude, the latter outcome by not using extremely attractive blue vane traps (Gibbs et al. 2017). Second, our estimates of local and landscape floral area assumed flower production is solely a function of the composition of plant communities of a given locale. In quantifying landscape floral area, we assumed plant communities of the same habitat type had equal numbers of flowers. In reality, the number, size, and quality of floral resources is influenced by many factors, including temperature, moisture availability, and soil fertility (Cardoza et al. 2012; Mu et al. 2015). For each site, we estimated local floral area based on relative abundance of plants present at the site but did not measure realized flowering in each year of our study. We suspect that soil fertility influenced bee abundance and richness at our study sites by changing flower production per unit vegetative cover, but lacking surveys of realized floral area at every site, we could not test this hypothesis. Third, we conducted bee surveys 1-3 years after Iverson et al plant surveys. Comparing habitat types, we expect the major trends in plant community composition, floral area, and richness (Figures S2-S4) would be robust to this 1-3 year time lag. In mesic upland remnant forests, floodplain forests, and forest edges, most species are long-lived trees and shrubs, so we expect little change in plant composition and floral area. In old fields, roadside ditches, mixed vegetable farms, and apple orchards, mowing, spraying, or crop rotation may have shifted plant communities in the time between plant and bee sampling. From our field observations, none of our sites experienced major disturbances like fire or pest outbreak. Fourth, by choosing to analyze our data with random forest models, we could not assess interactions between local and landscape quality. Local and landscape quality metrics were only weakly correlated across scales (minimum, mean, and maximum Pearson’s *r* = -0.6, 0.01, and 0.56, respectively, Figure S10), potentially suggesting independent effects of local and landscape quality, but we could not examine these questions. Finally, while assessing conservation plantings for pollinators, it has been a persistent challenge to assess whether adding flowers for bees increases wild-bee populations, or just concentrates individuals in high-quality patches. Our multi-dimensional characterization of local quality cannot definitively show that high-quality sites improved bee fitness, compared with drawing in more individuals that were already present in the landscape. In the future, to provide a measure of wild-bee reproductive output, we recommend researchers use passive traps or visual observation in tandem with nest boxes or colonies of managed bees (see Martínez-Núñez et al. 2020; Centrella et al. 2020). Two recent studies also presented promising methods to detect landscape-scale changes in wild-bee abundance or richness (Kleijn et al. 2018; Scherber et al. 2019), and it would be valuable to test these methods in more complex landscapes, such as in our study region.

## 5. Conclusion

In this study, we identified key components of landscape and local quality for wild bees, generating essential targets for bee conservation programs. Building on previous research showing that bees respond to both landscape and local resources, we found that the most relevant spatial scale varies by season and can differ when considering bee abundance or richness. Spring bees were heavily dependent on landscape resources, but abundance of summer bees responded to local, rather than landscape quality, implying that site-level management is most likely to be successful in supporting (or at least attracting) summer bees. In highly agricultural landscapes, however, improving local quality for summer bees will likely primarily benefit disturbance-tolerant, generalist species because rare bees are typically absent from these landscapes (Harrison et al. 2019). By quantifying landscape-scale floral resources, we found that semi-natural habitats, specifically forests, provision spring bees, likely by providing large pulses of resources from flowering trees. Many areas in our study region already have large patches of forest habitat, suggesting that adding more trees or shrubs represents a small marginal gain when compared with existing resources. Rather, to support spring bees in landscapes with relatively high forest cover, we recommend targeted additions or restorations of wetlands and surface-water features, habitats that should also provide conservation benefits for other plant and animal taxa. By considering spatial and temporal variation in resources, we developed context and season-specific recommendations to improve habitat quality for wild bees, a critical component of conservation efforts to offset the manifold stressors threatening these taxa.

## 6. Acknowledgments

We thank the many private landowners, Cornell Botanic Gardens, and Finger Lakes Land Trust who generously allowed us to collect bees on their property. We are grateful to Bryce Buck, Cheyenne Dolinsky, Ryan Reynolds, Liz Wagner, Darian Kraft, Kathy Wholaver, and Ginamaría Román Echevarría for assistance with field work and processing bee specimens, and logistical support from Kate Anton. We greatly appreciate assistance from Dr. David Biddinger and Dr. Robert Jean in identifying the bees we collected. Lastly, we thank Sarah Goslee, Dave Mortensen, and members of the Grozinger lab for feedback on early versions of this manuscript.

## Statements and Declarations

### Funding

This project was funded by a United States Department of Agriculture National Institute of Food and Agriculture (USDA-NIFA) pre-doctoral fellowship PENW-2017-07007 to MK, grant #2021-67021-34146 to CMG, and a graduate student grant from the USDA NIFA Northeast Sustainable Agriculture Research and Education program under subaward GNE17-142. We were also supported by the Foundation for Food and Agriculture Research grant #549032 to CMG and the College of Agricultural Sciences at Penn State USDA NIFA and Hatch Appropriations under Project #PEN04606 and Accession #1009362. We are grateful for support from the Pennsylvania State University Center for Pollinator Research Apes Valentes grant, College of Agricultural Sciences graduate student grant, and Intercollege Graduate Degree Program in Ecology.

This research was supported in part by the U.S. Department of Agriculture, Agricultural Research Service. The findings and conclusions in this manuscript are those of the author(s) and should not be construed to represent any official USDA or U.S. Government determination or policy. Mention of trade names or commercial products in this publication is solely for the purpose of providing specific information and does not imply recommendation or endorsement by the USDA. USDA is an equal opportunity provider and employer.

## Competing Interests

The authors have no relevant financial or non-financial interests to disclose.

## Author Contributions

MK, JFT, and CMG conceived and designed the study. KL and ALI created high-resolution land-use data, ALI collected plant community composition, floral density, and flower size datasets, and MK conducted soil and wild-bee sampling. MK analyzed the data and led manuscript writing. All authors contributed critically to the drafts and gave final approval for publication.

## Data Availability

All data required for the analyses presented here will be permanently archived in the Dryad Digital Repository. Code necessary to complete the analyses described here are archived on Zenodo (Kammerer 2021).

## Notes

### Competing Interest Statement

The authors have declared no competing interest.

### Summary of Updates

Correct typo in title.

