## Supplemental Information for "Seasonal bee communities vary in their responses to local and landscape scales: implication for land managers"

Figure S1: Non-metric multidimensional scaling of plant communities at 144 sites in the Finger Lakes region of New York USA (Iverson et al., in review). Sites are shown with colored points and polygon outlines differentiate 21 habitat types (Panel A) represented in the dataset. Plant species most associated with each habitat type (highest axis loadings) are labeled in black text, with all species represented with grey points (Panel B). stress= 0.153

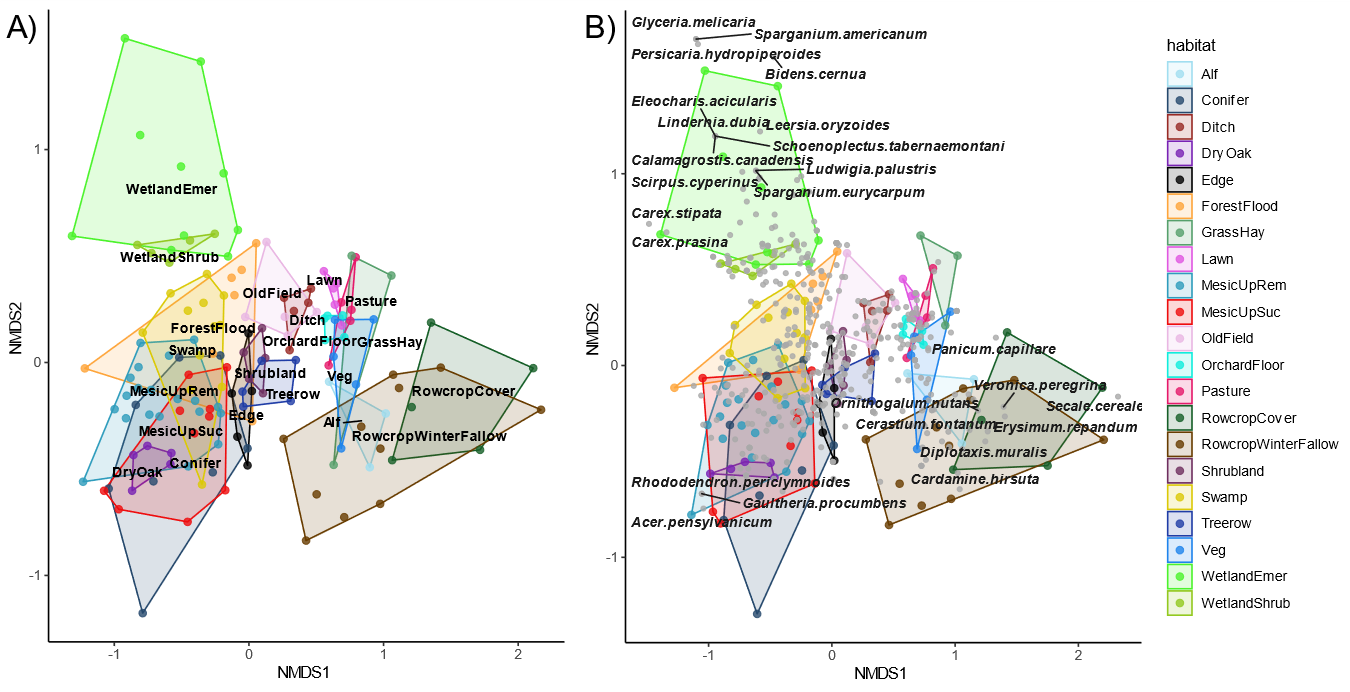

Figure S2: Ordination plots from principal components analysis of soil characteristics at 33 sites in the Finger Lakes region of New York USA. In panel A, site-centroids are shown with grey points and soil properties in burgundy vectors.

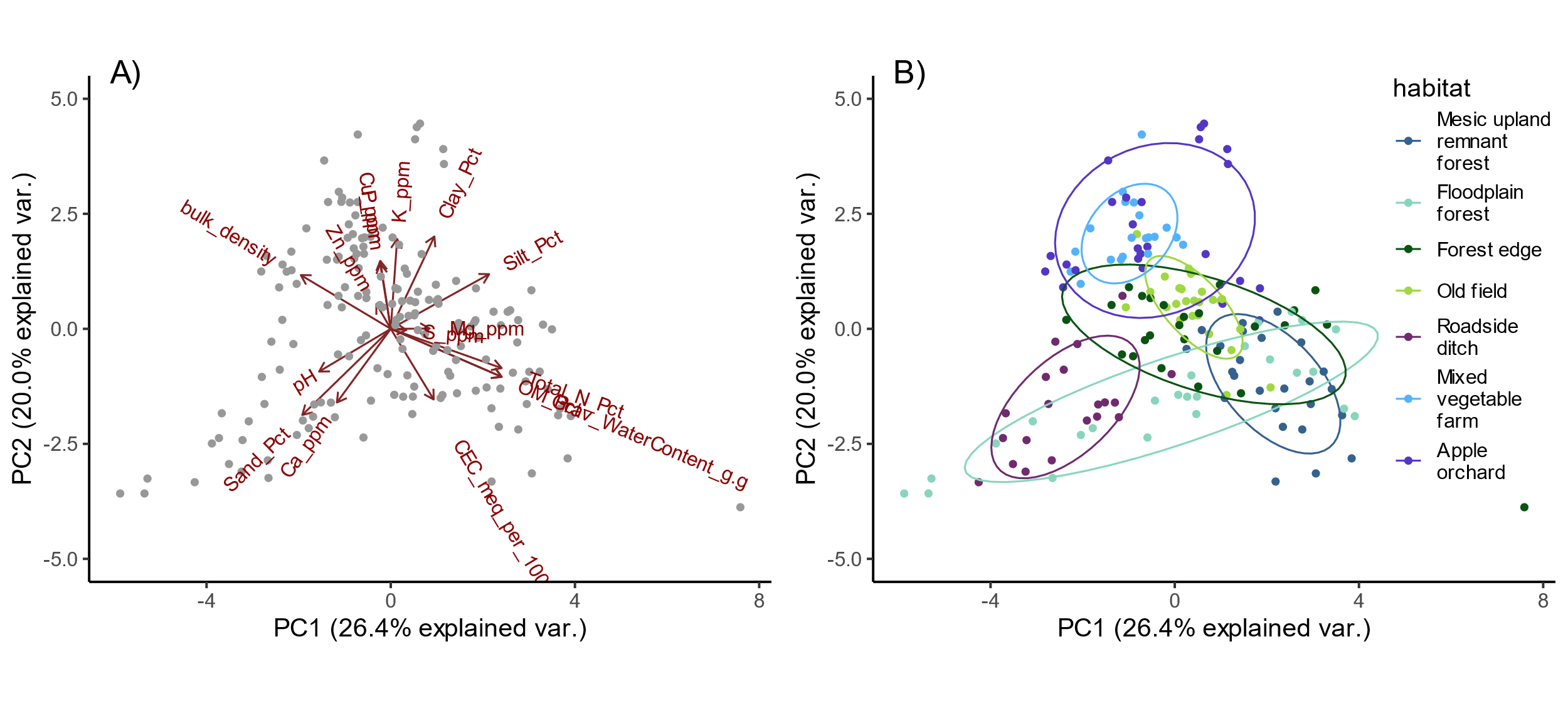

Figure S3: Floral area over time available in seven habitats types in the Finger Lakes region of New York, USA. Floral area values represent only insect-pollinated plants.
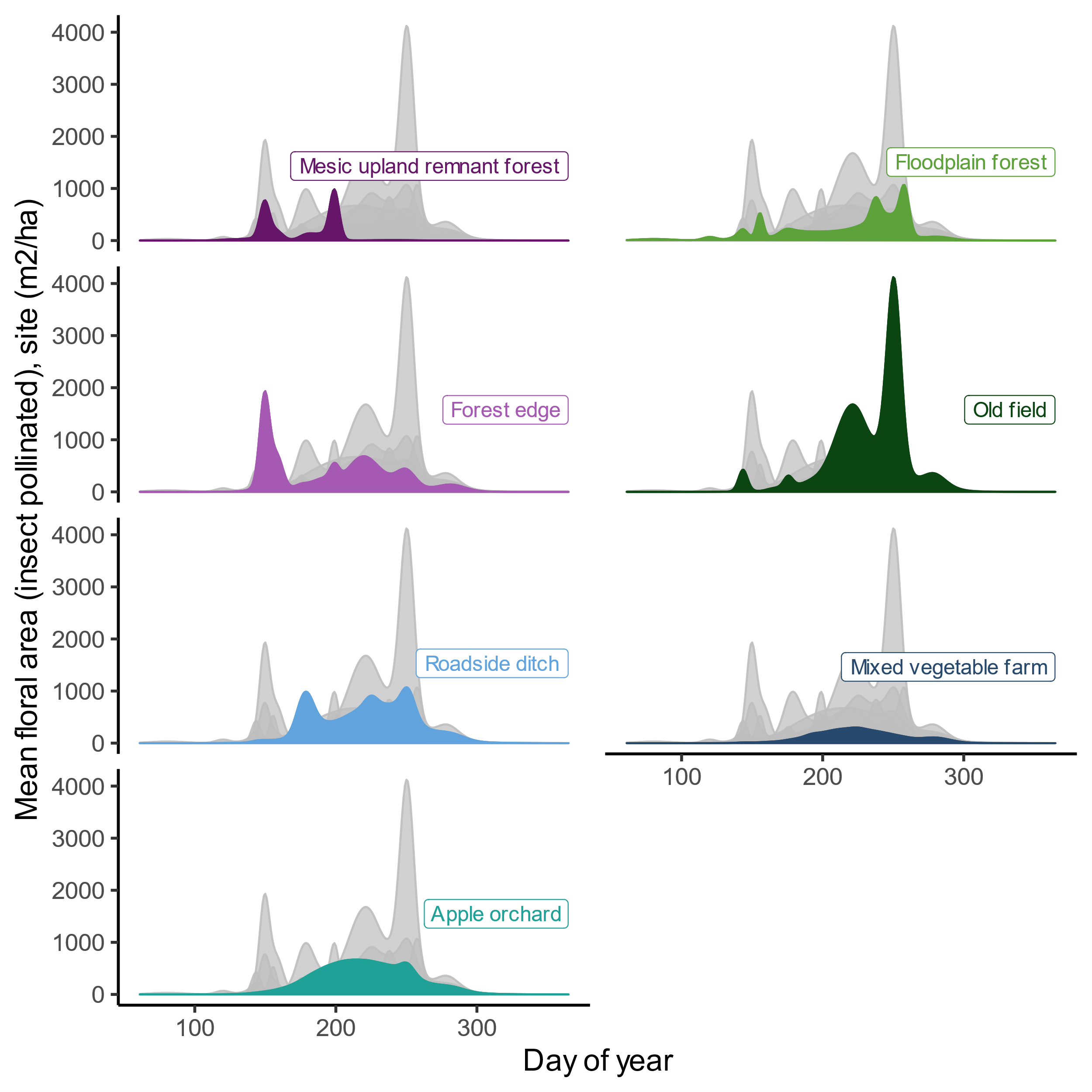

Figure S4: Species richness of all plants and insect-pollinated plants at 33 sites in the Finger Lakes region of New York, USA. Habitat types are abbreviated as follows: Apple = apple orchard , Ditch = roadside ditch, Edge = forest edge, Field = old field, Flood = floodplain forest, Forest = mesic upland remnant forest, Veg = mixed vegetable farm.

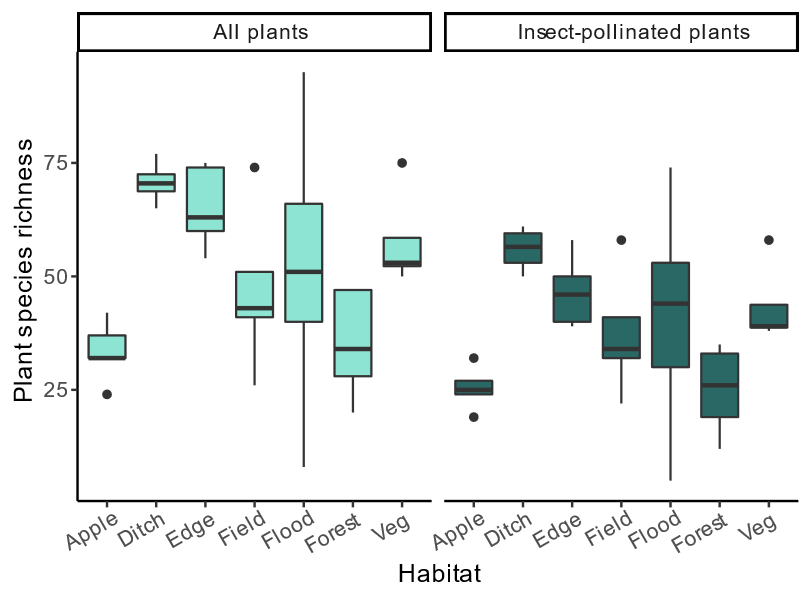

Figure S5: Composition of landscapes surrounding 33 sites in the Finger Lakes region of New York, USA. Data are shown as density overlaying frequency (histogram) plot.
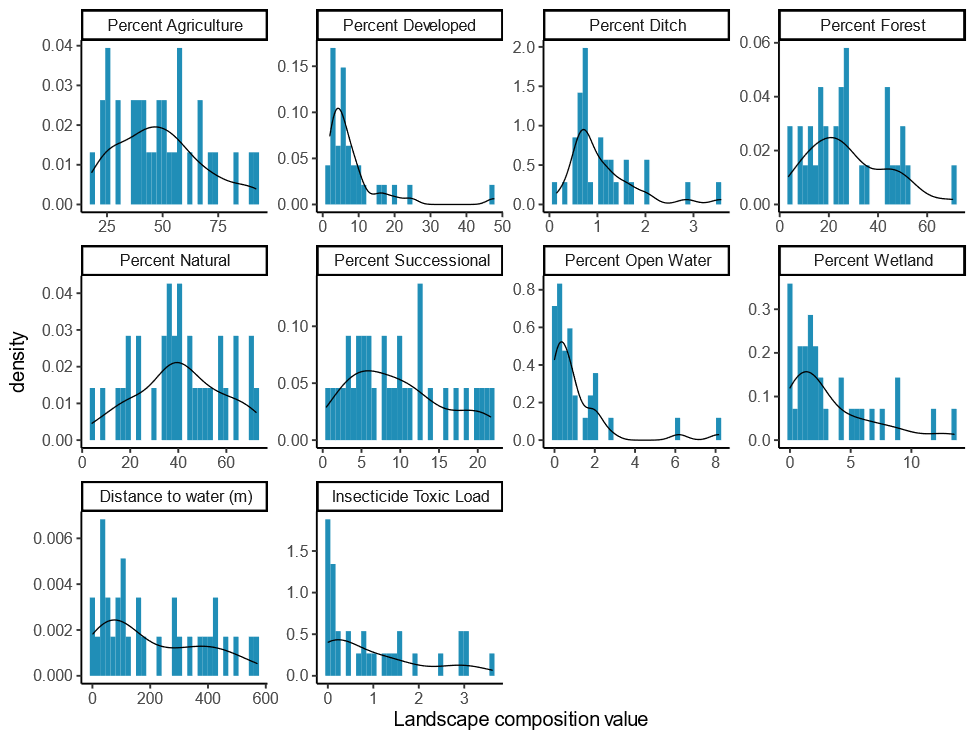

Figure S6: Topographic variables from 33 sites in the Finger Lakes region of New York, USA. Data are shown as density overlaying frequency (histogram) plot.

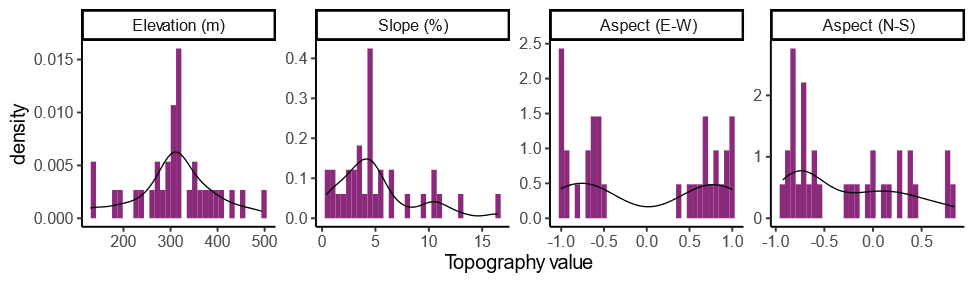

Figure S7: Configuration of landscapes surrounding 33 sites in the Finger Lakes region of New York, USA. Data are shown as density overlaying frequency (histogram) plot. Abbreviations are as follows: ed= edge density, enn_cv = coefficient of variation of euclidean nearest-neighbor distance, iji = interspersion and juxtaposition index, para_mn = mean perimeter-area ratio, shdi = Shannon's diversity index, sidi = Simpson's diversity index (Hesselbarth et al., 2019).

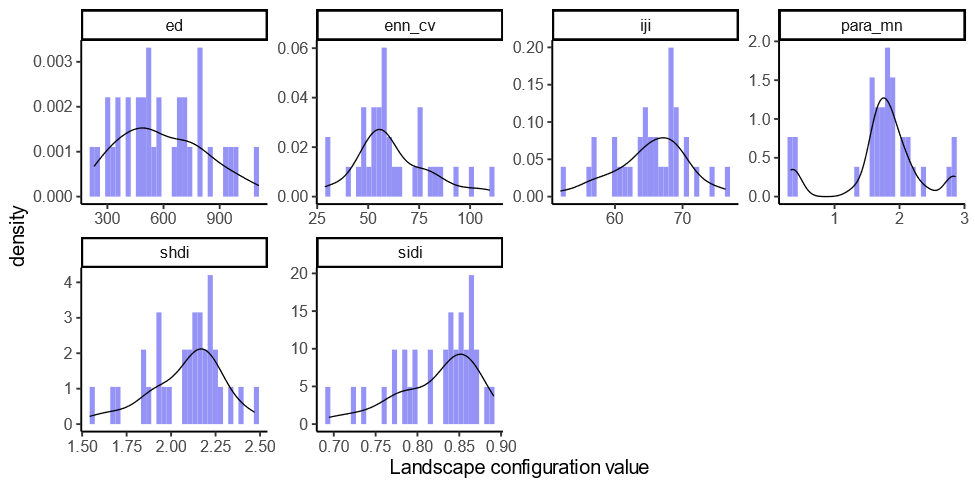

Figure S8: Abundance (panel A) and richness (B) of wild bees in spring and summer at 33 sites in the Finger Lakes of New York, USA. Species richness presented here was adjusted with coverage-based rarefaction, to account for variation in sampling effort. Habitat types are abbreviated as follows Apple = apple orchard , Ditch = roadside ditch, Edge = forest edge, Field = old field, Flood = floodplain forest, Forest = mesic upland remnant forest, Veg = mixed vegetable farm.

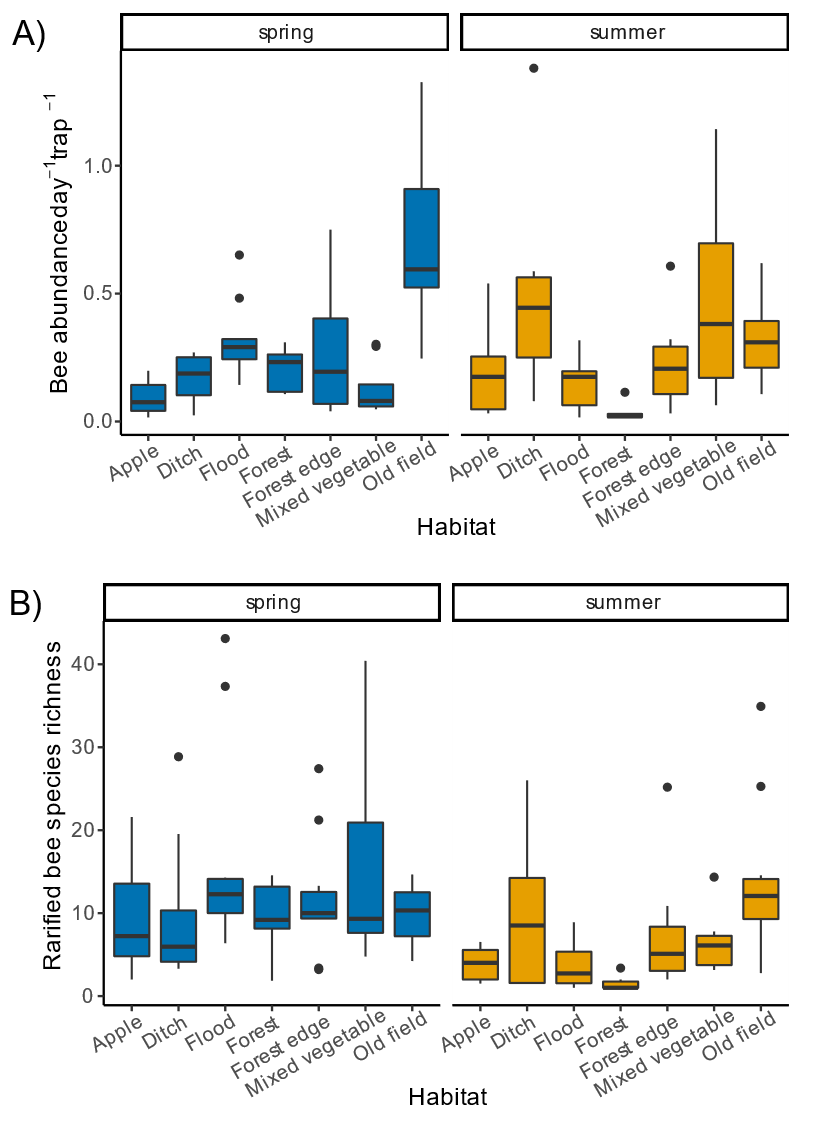

Figure S9: Relationship between landscape and site predictors and wild-bee abundance and richness in the spring. Here we show #5-7 most important predictors, with x-axis truncated to 10-90% quantiles (n=29 sites). To enable comparing relative effects across seasons with varying mean abundance or richness (Table 1), we depict values on the y-axis as difference from the predicted mean.

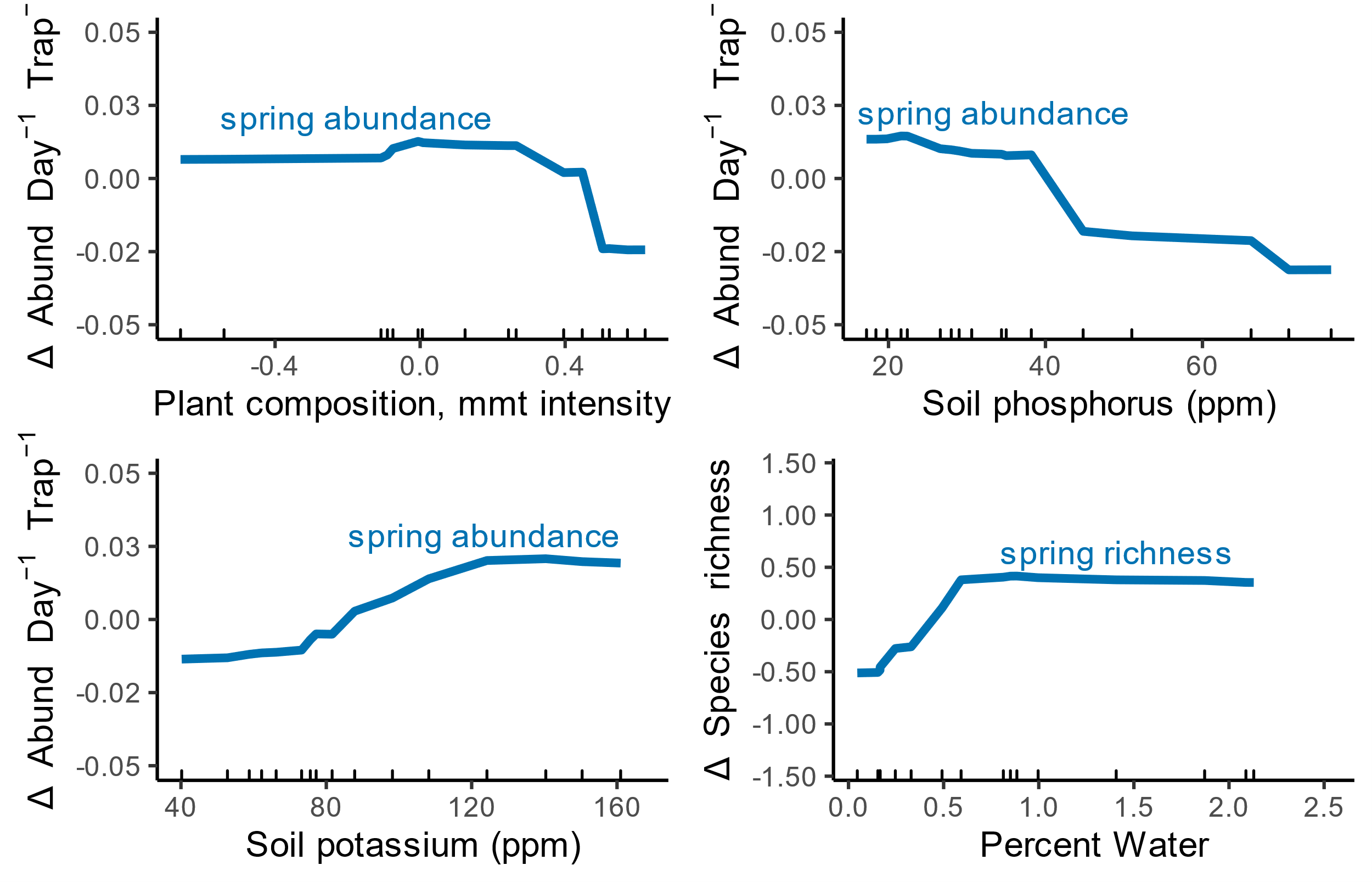

Figure S10: Coefficients from pairwise Pearson correlations of all site and landscape variables (n=72) included in random forest models. The black box highlights correlation coefficients from pairs that included one site variable and one landscape variable. ‘.IP’ indicates an insect-pollinated plant community, while ‘.all’ denotes all plants species

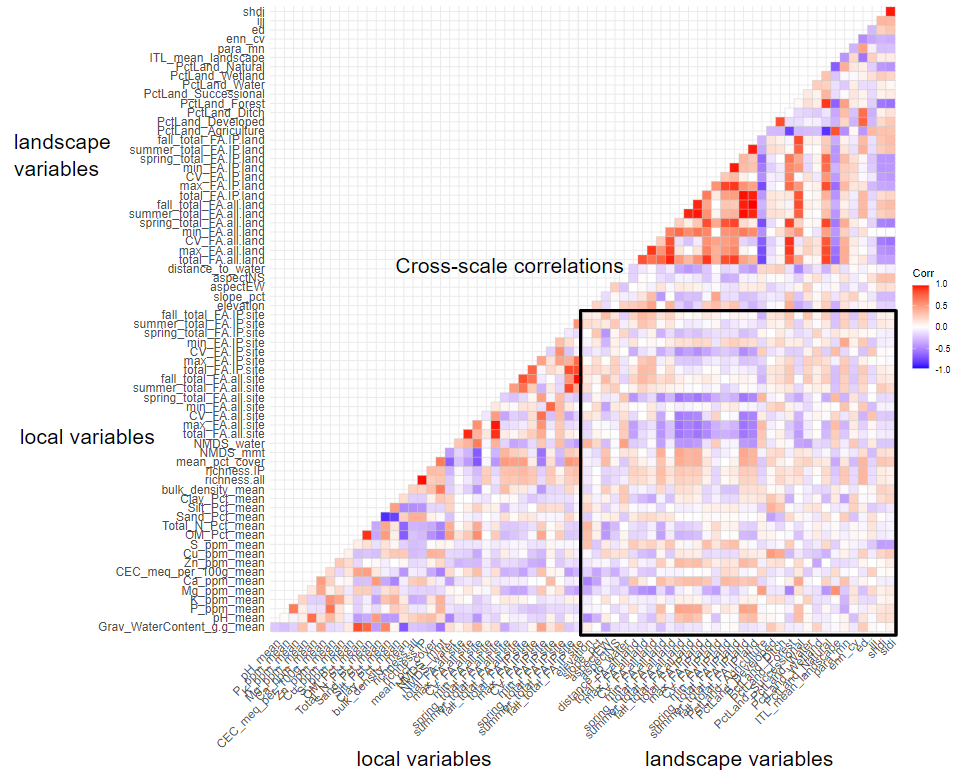

Table S1: Classification of land-cover classes into agriculture, forest, successional, wetlands, water, and developed habitats. Land-cover classes listed here are from high-resolution spatial data used to calculate landscape composition and configuration (Iverson et al, in review).

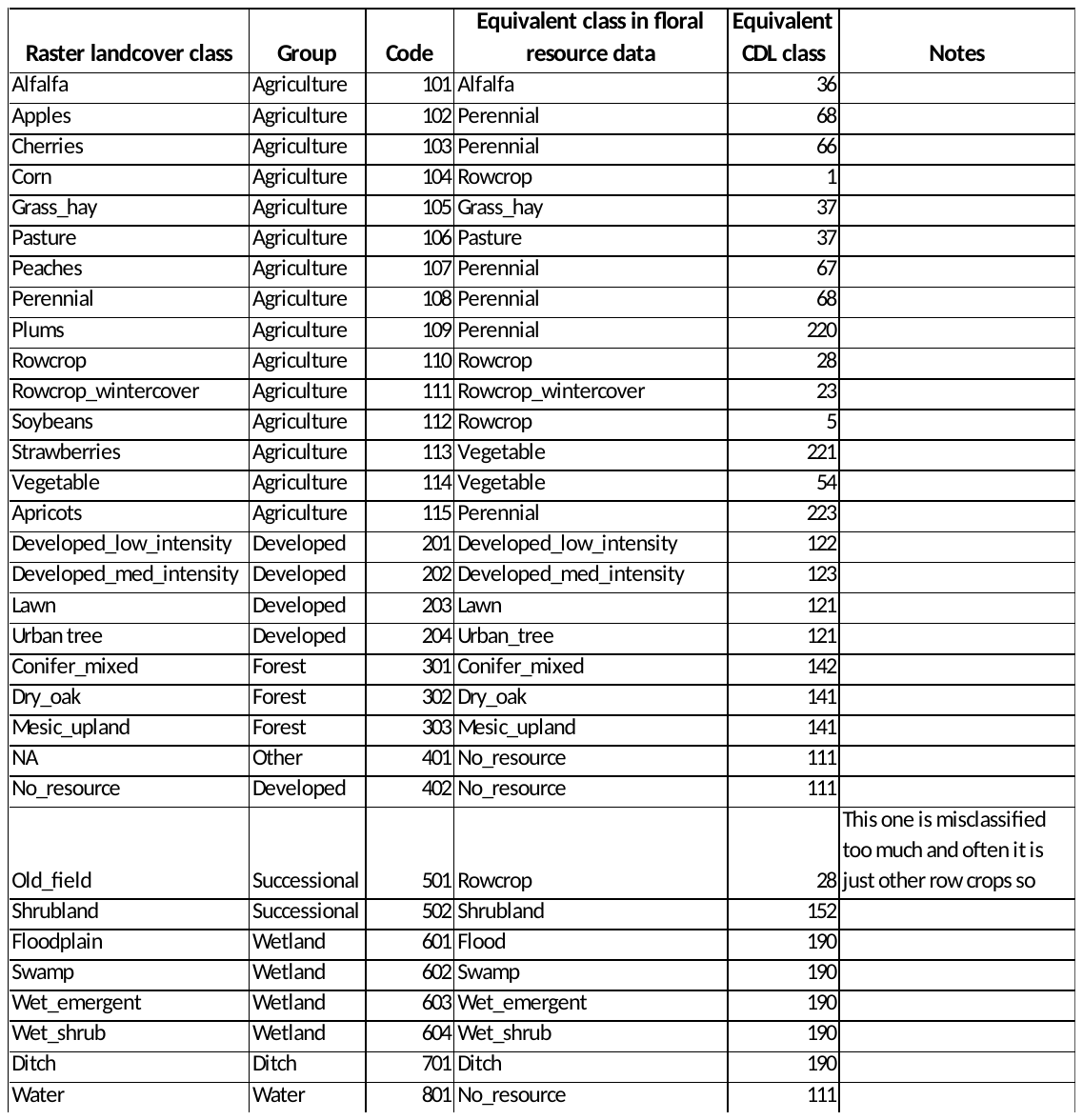
